# Distinct prefrontal-amygdala connectivity drives consolidated fear memories

**DOI:** 10.64898/2026.08.14.744891

**Authors:** Matthew Kenna, James Kesby, Li Xu, Robert Sullivan, Roger Marek, Pankaj Sah

## Abstract

Elucidating the neuronal circuitry that underpins memory formation is critical to understanding how organisms use past experience to guide adaptive behaviour. While memory formation has long been framed as the reactivation of a static ensemble of neurons established during initial learning, growing evidence suggests that memory traces are highly dynamic and undergo substantial reorganisation during consolidation. During the formation of auditory fear memory, initial acquisition is primarily mediated by the basolateral amygdala (BLA), whereas long-term expression relies on the medial prefrontal cortex (mPFC). However, the circuit motifs that coordinate this systemic redistribution remain poorly understood. Here, using targeted anatomical tracing and electrophysiology, we show that the reciprocal connectivity between the mPFC and BLA is organised as a parallel topography along the rostro-caudal axis. Leveraging this novel anatomical understanding of reciprocal communication between the amygdala and prefrontal cortex, we reveal an underlying circuitry mechanism by which fear memory traces are redistributed into subcortical-cortical networks after learning. Using activity-dependent engram capture and optogenetic manipulation, we demonstrate that post-learning engagement of a distinct sub-circuit linking the rostral BLA and rostral mPFC is a hallmark of the consolidated fear memory. These insights reveal that the consolidated engram requires the targeted engagement of a post-learning engram circuit, rather than a simple reactivation of neurons engaged during initial learning.

## Introduction

An overarching goal of neuroscience is to understand how the brain forms, stores, and retrieves memories. Memory formation is a fundamental adaptive process necessary for survival, enabling organisms to map threats and navigate changing environments. While standard neurobiological models divide memory into distinct phases of acquisition, consolidation, and retrieval - consolidation has been viewed as the passive solidification of cellular changes established during initial learning^1^. However, recent evidence indicates that memory consolidation is a dynamic process involving the active redistribution of the memory trace from localised, subcortical hubs to distributed cortical networks^2–6^. Identifying the precise circuit mechanisms that drive these post-learning alterations remains a major challenge, particularly as sensory cues are absent during consolidation to stimulate network activity.

The neural basis of memory has long centred on the engram; the idea that learning induces lasting physiological changes in a specific subset of neurons, forming a connected ensemble that is reactivated upon retrieval of the memory^7–9^. The application of immediate early gene (IEG)-based engram tagging techniques has confirmed that stimulation of learning-induced ensembles can artificially drive expression of the original memory^5,10–13^. However, these approaches routinely reveal a low cellular overlap (often 5–30%) between cells active during initial learning and those engaged during retrieval^14–17^. It remains unclear whether this modest overlap reflects the capture of non-engram-related activity (e.g. session-specific sensory features or movement-responsive cells) or dynamic shifts in the underlying memory ensemble between learning and retrieval.

The anatomical and functional connectivity between the BLA and mPFC provides an ideal framework to test this dynamic engram hypothesis. The initial acquisition of auditory fear memories is primarily mediated by the BLA, whereas long-term memory retrieval and expression critically depend on the mPFC^18–23^. How the memory trace scales from a subcortical structure during learning to a redistributed network that includes the mPFC during retrieval is unknown. While electrophysiological recordings from the PL during fear conditioning consistently show changes in neural activity following CS-US pairings, reported response profiles vary across studies^24–26^. These discrepancies may stem, in part, from the assumption that prefrontal subdivisions, such as the PL, are functionally and structurally homogeneous across the rostro-caudal axis^27–29^. In the mouse brain, the PL spans over 1.6 mm along the rostro-caudal plane, shifting significantly in its dorso-ventral position and local microcircuitry^27,30^. Examining BLA–mPFC communication and neural activity during fear memory formation along this domain could help elucidate the circuit mechanisms that govern how a memory scales over time.

To address these questions, we investigated changes in active neuronal ensembles across the full rostro-caudal and laminar axes of the mPFC before and after memory consolidation. By combining high-resolution circuit mapping with optogenetics, we demonstrate that a parallel sub-circuit linking the rostral BLA to the rostral mPFC undergoes selective post-learning engagement, serving as a critical substrate for consolidated memory expression.

## Results

### Retrieval of consolidated fear memory evokes layer 2/3 activity in the mPFC

Previous investigations of the neural circuits that underpin auditory fear memory have shown that the mPFC is recruited following the initial learning phase and is required for memory retrieval^21–24,26,31^. To understand the changes in neural activity across the pre-and post-consolidation windows that may reflect this engagement, we applied activity-dependent engram tagging within this circuit framework. Specifically, to reveal the neural correlates of fear memory formation across the mPFC, we virally delivered the Robust Activity Marking (RAM) system^32^ to drive expression of the red fluorophore mScarlet in active neuronal ensembles during precise behavioural windows. We compared active prefrontal networks across four distinct behavioural cohorts: a home cage control group, a tone-only control group, a fear learning group tagged during Pavlovian auditory fear conditioning, and a fear retrieval group tagged during memory retrieval, two days post-conditioning (see Methods; **Fig. 1a, c, d**). Behavioural tracking confirmed robust and comparable fear memory acquisition in the conditioned cohorts, while tone-only presentations elicited no significant changes in freezing (**Fig. 1b**). Robust freezing responses were observed in the retrieval cohort during the subsequent memory retrieval test, indicating that this captured neuronal ensemble is associated with the expression of a robust fear memory (**Fig. 1b**).

**Figure 1.**
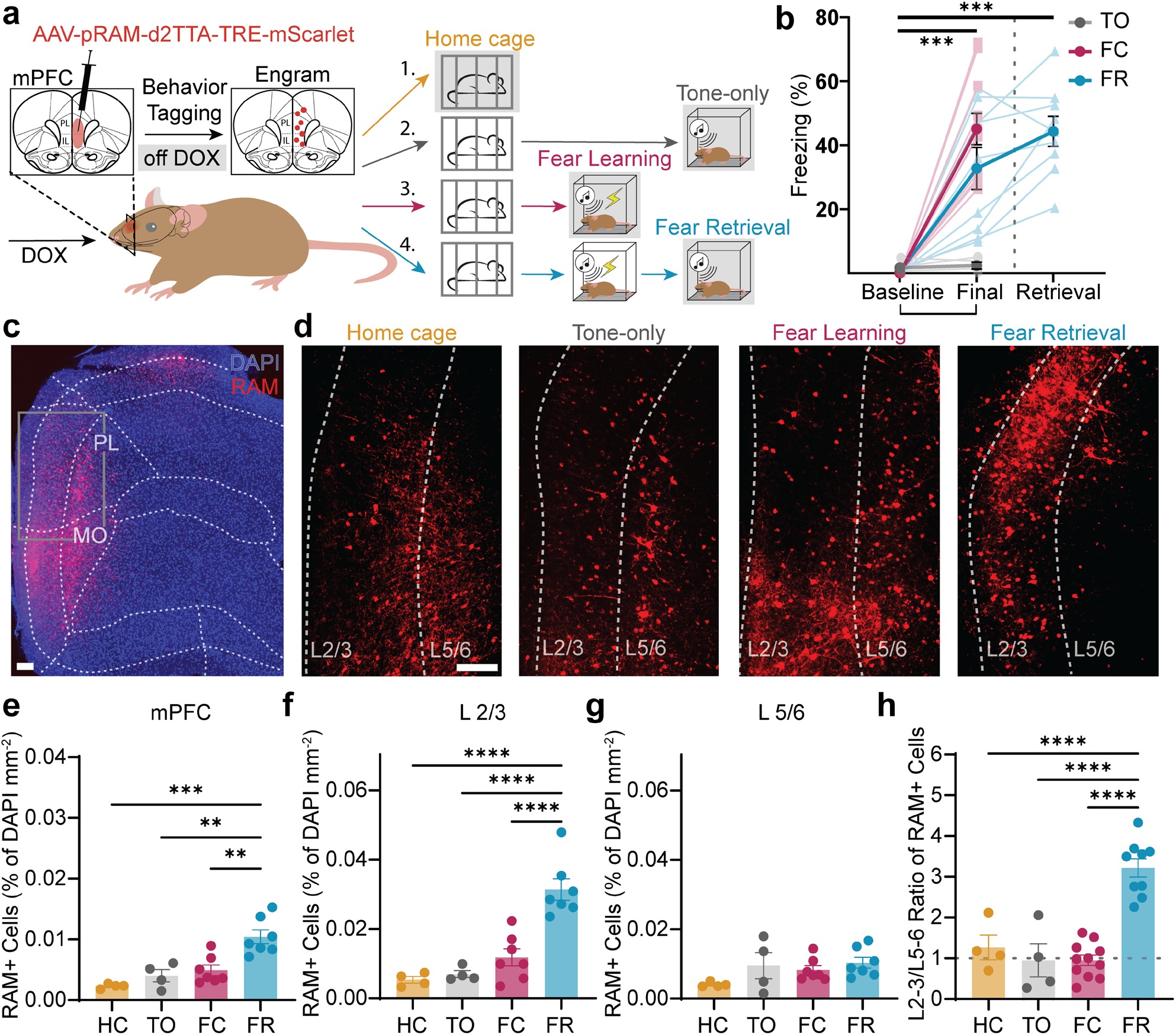
Retrieval of consolidated fear memory engages layer 2/3 activity across the mPFC. **a.** Experimental configuration detailing the viral injection of the activity-dependent RAM system across the rostro-caudal extent of the mPFC to express mScarlet in neurons active during restricted, Dox-free behavioural windows. Four experimental cohorts were tested: home cage control (HC, yellow, n = 4); tone-only control (TO, grey, n = 4); fear learning (FC, purple, n = 11); and fear retrieval (FR, blue, n = 9). Grey shaded area indicates behavioural window when doxycycline was removed from the feed and cells tagged by the RAM system. **b.** Freezing scores across behavioural sessions. Tone-only presentations (grey) did not significantly increase freezing behaviour (baseline vs. final: P = 0.70). Auditory fear conditioning (purple) drove significant increases in freezing across trials in both the fear learning group (baseline vs. final: P < 0.001) and the fear retrieval (blue) group (baseline vs. final: P < 0.01, two-way ANOVA with Tukey’s multiple comparisons test). Presentation of the CS during the retrieval test drove significant freezing relative to baseline (P < 0.001). **c.** Example confocal image of RAM-labelled (mScarlet+) neurons within the mPFC in the tone-only condition. Grey rectangle indicates the location of the area depicted in **d**. Scale bar = 100 µm. **d.** Representative high-magnification confocal images of RAM-labelled somata within superficial (L2/3) and deep (L5/6) cortical layers across the four behavioural groups. Scale bar = 100 µm. A/P coordinates: HC = +2.4 mm Bregma; TO = +2.5 mm Bregma, FC = +2.4 mm Bregma, FR = +2.6 mm Bregma. **e.** Mean number of active RAM-labelled cells per brain slice (% of DAPI/mm^2^) across the mPFC. Fear retrieval induced a significant increase in total active cell count relative to all other groups (P < 0.01, one-way ANOVA with Tukey’s test). **f.** Mean number of active RAM-labelled cells per slice in L2/3 (% of DAPI/mm^2^). Fear retrieval drove a significant increase in superficial cell recruitment relative to all other conditions (P < 0.0001, one-way ANOVA with Tukey’s test). **g.** Mean number of active RAM-labelled cells per slice in L5/6 (% of DAPI/mm^2^), showing no significant differences across cohorts (P > 0.05, one-way ANOVA). **h.** Superficial-to-deep ratio (L2/3 vs. L5/6) of active cells across the mPFC. Fear retrieval drove a significant increase in this ratio compared to home cage, tone-only, and fear learning groups (P < 0.0001, one-way ANOVA with Tukey’s test). All data are presented as mean ± SEM.

Quantification of RAM-labelled cells across the mPFC revealed that following memory consolidation fear retrieval produced a significant increase in the number of active cells, compared to all other conditions (**Fig. 1e**, and Extended Data Fig. 1). This post-consolidation increase was restricted to superficial layers; fear retrieval drove a substantial increase in active cells within layer 2/3 (L2/3) relative to home cage, tone-only, and fear learning groups (**Fig. 1f**), while the number of active cells within deep layers (L5/6) was unchanged across all behavioural states (**Fig. 1g**). Consequently, there was a significant increase in the ratio of active cells in L2/3 to L5/6 exclusively during fear retrieval compared to all other groups (**Fig. 1h**). To validate this result using a different marker of activity, we examined a cohort of RAM-mScarlet injected animals immunostained for c-Fos two hours after fear retrieval. These data confirmed a significant overlap between the RAM-labelled and c-Fos-expressing populations, but the RAM system captured a substantially larger population of inhibitory neurons, particularly within the superficial layers (Extended Data Fig. 2). This differential targeting may explain why traditional c-Fos labelling alone often underrepresents layer 2/3 activity and fails to capture this specific laminar pattern during retrieval.

These data show that retrieval of a consolidated auditory fear memory drives a distinct, layer-specific pattern of activity in the mPFC characterised by dominant L2/3 activation. Importantly, because a significant shift in the tone-evoked mPFC activity pattern occurs after the learning period, these findings could indicate a post-learning engagement of a specific prefrontal network.

### Fear memory retrieval engages activity in superficial layers of the rostral mPFC

Given the global shift toward superficial prefrontal activity during fear retrieval, we next examined its distribution along the anterior-posterior axis by partitioning the mPFC into rostral, intermediate, and caudal zones (see Methods; **Fig. 2a**). Mapping the coordinates of RAM-labelled neurons revealed a distinct topographical divergence (**Fig. 2b, c**). During fear learning, no significant differences were detected in the number of active cells between superficial and deep layers at any point along the rostro-caudal axis (**Fig. 2d**). In contrast, following consolidation, fear retrieval engaged a significantly greater number of active cells within L2/3 of the rostral mPFC compared to deep layers (**Fig. 2d**). Notably, while a broad rostral-to-caudal gradient of decreasing L2/3 activation was apparent across the mPFC, no statistically significant layer-specific differences were found within the intermediate or caudal zones during retrieval.

**Figure 2.**
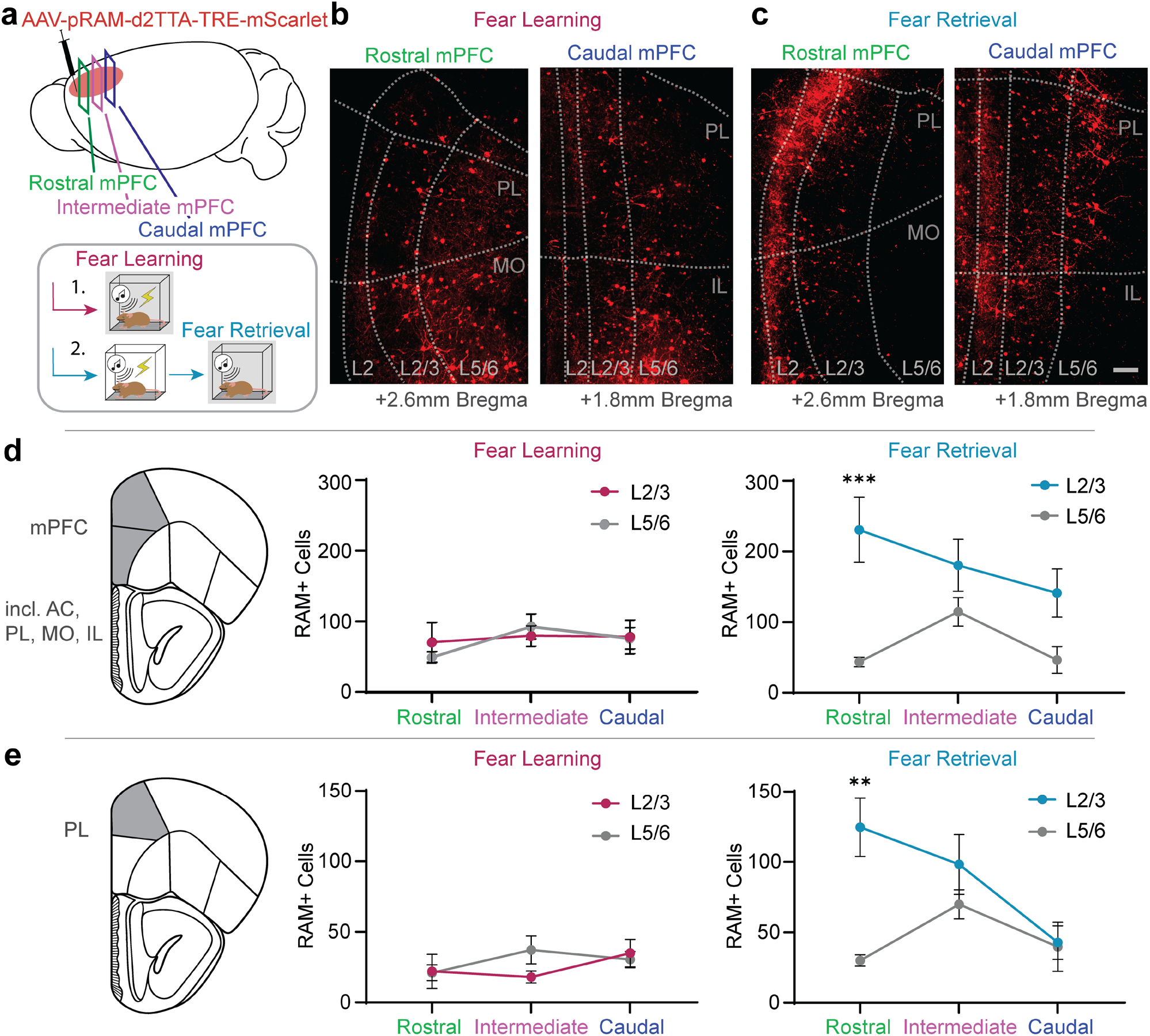
Post-consolidation increase in superficial activity is driven by the rostral mPFC and prelimbic cortex. **a.** Spatial schematic of the analysis strategy. The mPFC was partitioned into rostral (green), intermediate (magenta), and caudal (dark blue) zones along the anterior-posterior axis to evaluate RAM-labelled networks before (fear learning, purple) and after consolidation (fear retrieval, light blue, 48 hr post-learning). Shaded regions show behavioural states where animals were off DOX and RAM system engaged. **b.** Representative confocal images of RAM-labelled ensembles during fear learning in the rostral (left) and caudal (right) mPFC planes. Scale bar = 100 µm. PL = prelimbic cortex; MO = medial orbital cortex; IL = infralimbic cortex. **c.** Representative confocal images of RAM-labelled ensembles during fear retrieval in the rostral (left) and caudal (right) mPFC planes. Scale bar = 100 µm. **d.** Total number of active cells within L2/3 and L5/6 across the rostral, intermediate, and caudal zones of the entire mPFC labelled during fear learning (middle) and fear retrieval (right). Distribution of fear learning cells showed no significant differences between layers along the rostro-caudal axis (P > 0.05, two-way ANOVA with Šídák’s post-hoc test). Conversely, fear retrieval drove a significant increase in active cells within L2/3 of the rostral mPFC relative to corresponding deep layers (P < 0.01, two-way ANOVA with Šídák’s test). No significant layer-specific differences were found in intermediate or caudal segments (P > 0.05). AC = anterior cingulate cortex. **e.** Layer-specific count of active cells restricted within the prelimbic (PL) cortex labelled during fear learning (left panel) and fear retrieval (right panel). Distribution of fear learning cells showed no layer-specific differences across the rostro-caudal axis (P > 0.05, two-way ANOVA). Fear retrieval produced a significant increase in active cells within L2/3 of the rostral PL compared to its deep layers (P < 0.001, two-way ANOVA with Šídák’s test). All data are presented as mean ± SEM.

Given the well-documented role of the prelimbic (PL) cortex in driving fear memory retrieval^19,21,22^, we next restricted our spatial analysis to the PL, which uniquely spans the entire longitudinal extent of the mPFC. Neuronal ensembles captured during fear learning exhibited a uniform distribution across both superficial and deep layers throughout the rostro-caudal axis of the PL (**Fig. 2e**). Following consolidation, fear retrieval drove a highly localised and significant increase in active cells within L2/3 of the rostral PL, relative to L5/6 (**Fig. 2e**). Again, we observed a descending rostro-caudal trend in superficial layer activity across the longitudinal axis; although L2/3 active cells were enhanced within both the intermediate and caudal PL, these did not differ significantly from those found in corresponding deeper layers. Thus, the post-learning increase in active cells in superficial layers is not uniform across the mPFC, but is predominately restricted to the rostral subdivision of the PL.

This superficial layer dominance proved durable over time, remaining significantly elevated during a remote 7-day retrieval test (Extended Data Fig. 3). Furthermore, tagging ensembles following differential fear conditioning (see Methods) showed that this pattern of engagement across the rostro-caudal axis is stimulus-specific, as the threat-conditioned stimulus (CS+) drove a prominent rostral L2/3 elevation that was distinct to the pattern of activity of the unreinforced stimulus (CS−) (Extended Data Fig. 4). Finally, microinfusion of the NMDA receptor antagonist DL-2-amino-5-phosphonovaleric acid (APV, 50 mM) into the basolateral amygdala (BLA) immediately prior to fear conditioning impaired learning and selectively reversed the elevation of L2/3 active cells in the rostral PL during retrieval (Extended Data Fig. 5), showing that this localised prefrontal recruitment is dependent on fear memory formation.

These findings show that the post-consolidation increase in superficial prefrontal activity is driven principally by neurons in the rostral mPFC, specifically the rostral PL. This selective post-learning engagement indicates that the widespread assertion of functional homogeneity across the rostro-caudal axis of prefrontal subdivisions may be oversimplified. Instead, it suggests that engagement of circuits in the rostral mPFC occurs after the learning period, highlighting the need to examine the structural connectivity along the rostro-caudal axis to understand why these localised differences arise.

### Reciprocal connections between the mPFC and BLA are organised along the rostro-caudal axis

The mPFC has previously been implicated in driving the expression of fear memory through reciprocal connections with the BLA^22,33–35^. Anatomical and electrophysiological studies have established bidirectional, monosynaptic connections between the mPFC and BLA, with BLA afferents preferentially targeting superficial mPFC layers and reciprocal projections predominantly arising from layer 2/3 neurons^36–40^. Thus, the increase in superficial prefrontal activity observed during fear retrieval could reflect a post-learning engagement of this reciprocal circuit. However, given the distribution of active neurons during memory retrieval along the longitudinal plane, it remains unclear how these structures communicate across the rostro-caudal axis, as previous circuit models have assumed this connectivity to be uniform, particularly within sub-regions^36,41^.

To address this, we utilised a triple viral tracing approach to evaluate connectivity between the mPFC and BLA along the rostro-caudal axis. First, we injected anterograde viral tracers expressing distinct fluorophores into the rostral, intermediate, and caudal mPFC, ensuring that all injections encompassed the PL (**Fig. 3a, b** and Extended Data Fig. 6). Serial coronal sections were registered to the Allen Mouse Brain Atlas to measure brain-and area-normalised axonal fluorescence intensity along the longitudinal axis of the BLA, establishing a proportional distribution profile for each input pathway (see Methods; **Fig. 3c**). Total efferent fibre distribution to the entire BLA did not differ significantly among the three prefrontal zones (**Fig. 3d**). However, mapping fibre distribution profiles revealed a distinct topographical organisation: projections originating from the rostral mPFC terminated preferentially within the rostral BLA, whereas caudal mPFC afferents selectively targeted the caudal BLA (**Fig. 3e**).

**Figure 3.**
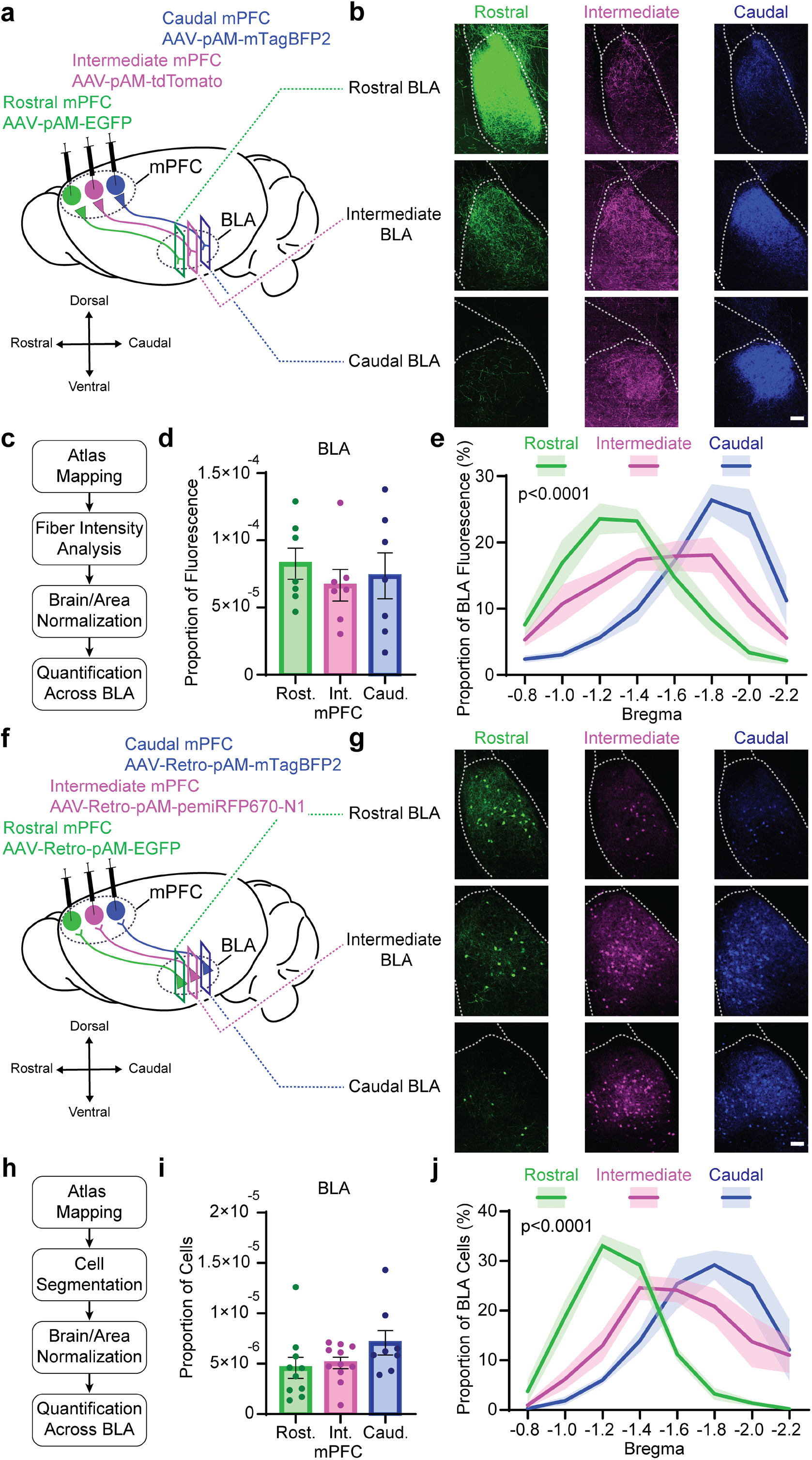
Prefrontal–amygdala connectivity is topographically organised along the rostro-caudal axis. **a.** Strategy for triple anterograde tracing. Viruses expressing distinct fluorophores were injected into the rostral (EGFP, green), intermediate (tdTomato, magenta), and caudal (mTagBFP2, blue) mPFC to trace efferent paths to the BLA. **b.** Example confocal images showing mPFC efferent fibres from the three prefrontal segments localising within the rostral, intermediate, and caudal BLA. Scale bar = 100 µm. **c.** Computational alignment and normalisation pipeline. Serial sections (200 µm) were mapped to the Allen Mouse Brain Atlas. Brain-and area-normalised fibre intensity values were calculated across the rostro-caudal extent of the BLA. **d.** Total area-normalised proportion of fibre input from the rostral, intermediate and caudal mPFC localised in the BLA. There was no significant variance in overall input magnitude among the three mPFC zones (P > 0.05, one-way ANOVA). **e.** Distribution profile mapping the proportion of total BLA fluorescence for each mPFC pathway along the rostro-caudal axis of the amygdala (n = 7 mice). The distributions along the rostro-caudal axis differed significantly across pathways (two-way repeated-measures ANOVA, pathway x bregma coordinate interaction, p < 0.0001). **f.** Strategy for triple retrograde tracing. Retrograde tracers expressing distinct fluorophores were delivered into the rostral (EGFP, green), intermediate (pemiRFP670, magenta), and caudal (mTAGBFP2, blue) mPFC. **g.** Representative confocal images of retrogradely labelled neurons within the rostral, intermediate, and caudal BLA targeting the three prefrontal zones. Scale bar = 100 µm. **h.** Cell segmentation and automated analysis pipeline mapping projection cell counts across atlas-aligned regions. Brain-and area-normalised cell counts were calculated across the rostro-caudal extent of the BLA. **i.** Total area-normalised proportion of projection cells targeting the rostral, intermediate, or caudal mPFC originating from the BLA, showing no significant differences in total projection magnitudes (P > 0.05, one-way ANOVA). **j.** Proportional distribution profile showing where prefrontal-projecting neurons were located along the rostro-caudal axis of the BLA (n = 12 mice). The distributions along the rostro-caudal axis differed significantly across pathways (two-way repeated-measures ANOVA, pathway x bregma coordinate interaction, p < 0.0001). All data are presented as mean ± SEM.

To determine if this parallel organisation is reciprocal, we injected distinct retrograde tracers into the same rostral, intermediate, and caudal prefrontal zones (**Fig. 3f, g**). Employing a similar atlas-registration pipeline, retrogradely labelled neurons were segmented and quantified across the BLA axis to determine the distribution of projection neurons for each target zone (**Fig. 3h**). The total proportion of BLA projection neurons targeting each prefrontal site was again equivalent, suggesting that the BLA input to each prefrontal region does not differ significantly (**Fig. 3i**). However, mapping these projection neurons along the axis of the BLA revealed a highly topographical pattern which mirrored the anterograde efferent distribution (**Fig. 3j**). Neurons projecting to the rostral mPFC were concentrated primarily within the rostral BLA, while those targeting the caudal mPFC were located within the caudal BLA. Crucially, restricting the rostral tracer exclusively to the medial orbital cortex (MO) yielded virtually no retrogradely labelled somata in the BLA (Extended Data Fig. 6d), demonstrating that projections specifically targeting the rostral PL drive the rostral portion of this topographical distribution.

Together, these anatomical tracing datasets reveal that reciprocal projections between the mPFC and BLA are topographically organised across the rostro-caudal axis as continuous spatial gradients. This shows that that the connectivity profiles of both the mPFC and BLA are not uniform across their anterior-posterior axes. Most notably, this arrangement reveals a reciprocal network linking the rostral mPFC and rostral BLA, a largely overlooked sub-circuit whose role in fear memory formation is yet to be examined.

### Synaptic connectivity between the BLA and mPFC is topographically organised along the rostro-caudal axis

To determine whether this parallel anatomical architecture corresponds to a functional organisation of synaptic connections, we designed an optical approach to independently stimulate two distinct synaptic inputs onto the same postsynaptic cell, termed independent Dual-Opsin Terminal Stimulation (iDOTS; **Fig. 4a**). Although red-shifted variants of channelrhodopsin have been engineered to permit independent optical control of neuronal activity^42^, the broad excitation spectrum’tail’ of these opsins causes unavoidable crosstalk when combined with conventional blue-light-activated tools, especially for larger post-synaptic responses. We addressed this limitation by combining the red-shifted opsin, Chrimson, with the inhibitory opsin, PdCO (Chr-PdCO)^43^, to allow for bidirectional optical control of a specific brain circuit. By expressing Chr-PdCO in one set of synapses and channelrhodopsin 2 (ChR2) in the other, we were able to independently recruit two distinct inputs while recording from a single postsynaptic cell without spectral crosstalk (see Methods; **Fig. 4b** and Extended Data Fig. 7). Applying this approach to resolve converging prefrontal inputs along the anterior-posterior axis, we virally expressed ChR2 in the rostral mPFC and Chr-PdCO in caudal mPFC. Then, acute *ex vivo* brain slices were prepared and whole-cell patch-clamp recordings obtained from BLA neurons across the longitudinal axis while recruiting synaptic inputs using 405nm and 640nm light (see Methods; **Fig. 4a**). Functional circuit mapping with iDOTS revealed that neurons within the rostral BLA received dominant monosynaptic inputs from the rostral mPFC, whereas caudal BLA neurons were driven primarily by the caudal mPFC (**Fig. 4c**). This topographical pattern matched the evoked excitatory postsynaptic current (EPSC) amplitudes, which were largest at targets corresponding to their parallel structural loops (**Fig. 4d**).

**Figure 4.**
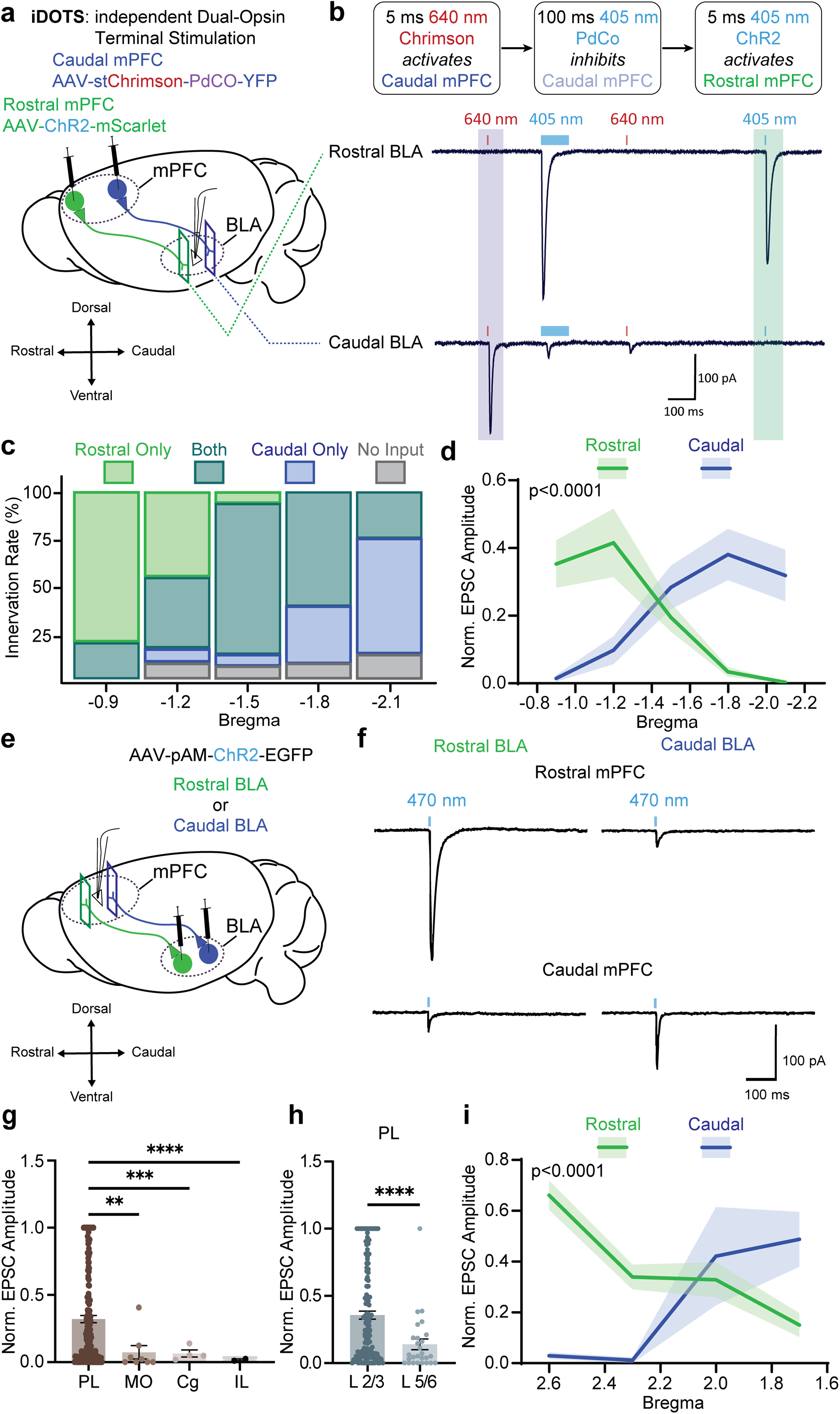
Synaptic connectivity between the mPFC and BLA is organised along the rostro-caudal axis. **a.** Schematic for the use of iDOTS to investigate projections from the rostral and caudal mPFC to the BLA. ChR2 was delivered into the rostral mPFC, and a viral vector of Chrimson and PdCO was delivered into the caudal mPFC. Whole-cell patch-clamp recordings were performed from neurons across the rostro-caudal axis of the BLA. **b.** Light delivery sequence for path-specific isolation. A brief (5 ms) 640 nm light pulse drives Chrimson to stimulate caudal mPFC inputs (blue shaded area). A subsequent long (100 ms) 405 nm pulse activates PdCO to shunt release from the same caudal terminals. A subsequent brief (5 ms) 405 nm light pulse then isolates inputs driven by ChR2 from the rostral mPFC (green shaded area). Below: representative EPSC traces recorded from individual neurons in the rostral and caudal BLA within the same animal. Shaded boxes designate the analysis windows for isolated inputs from the caudal (blue) and rostral (green) mPFC. All recordings were performed at –60 mV. **c.** Functional innervation rate across the rostro-caudal axis of the BLA, showing the percentage of recorded cells driven exclusively by the rostral mPFC (green), caudal mPFC (blue), convergent inputs (turquoise) or no input (grey). **d.** Within-animal normalised excitatory postsynaptic current (EPSC) amplitudes for rostral and caudal mPFC inputs across the rostro-caudal axis of the BLA. The distributions along the rostro-caudal axis differed significantly across pathways (two-way repeated-measures ANOVA, pathway x bregma coordinate interaction, p < 0.0001). **e.** Schematic of experimental approach to map the strength of synaptic input from the BLA to mPFC across the rostro-caudal axis. ChR2-EGFP was injected into the rostral (green) or caudal BLA (blue), and recordings were performed across the rostro-caudal axis of the mPFC. **f.** Representative EPSC traces recorded from rostral PL (+2.6 mm Bregma; top) and caudal PL neurons (+1.7 mm Bregma; bottom) during stimulation of inputs from the rostral (left) or caudal BLA (right). Blue bars indicate the 5 ms, 470 nm light pulse. All recordings were performed at –60 mV. **g.** Normalised EPSC amplitudes across prefrontal sub-regions. Neurons within the PL cortex received significantly larger inputs from the BLA compared to any other mPFC sub-region (P < 0.001, one-way ANOVA). **h.** Normalised EPSC amplitudes across cortical layers of the PL. Superficial L2/3 neurons received significantly larger BLA inputs compared to deep L5/6 neurons (P < 0.0001, unpaired t-test). **i.** Mapping of normalised EPSC amplitudes across the anterior-posterior axis of L2/3 PL neurons driven by rostral BLA (n = 111 neurons from 10 mice) or caudal BLA inputs (n = 21 neurons from 3 mice). The distributions along the rostro-caudal axis differed significantly across pathways (two-way repeated-measures ANOVA, pathway x bregma coordinate interaction, p < 0.0001). All data are presented as mean ± SEM.

We next mapped the functional connectivity in the reverse direction (BLA-to-mPFC) by delivering ChR2 into either the rostral or caudal BLA and performed whole-cell patch-clamp recordings across the prefrontal cortex (**Fig. 4e**). Synaptic input was highly sub-region specific, with the largest EPSCs recorded from principal neurons within the PL as compared to other prefrontal subdivisions (**Fig. 4g**). Within the PL, inputs displayed strict laminar selectivity, with significantly larger EPSCs in superficial L2/3 neurons than in deep L5/6 neurons (**Fig. 4h**). We therefore focused on L2/3 of the PL to evaluate the distribution of BLA input across the rostro-caudal axis. Due to this layer-subregion configuration spanning the entire length of the mPFC and receiving the majority of incoming amygdalar drive, it allowed us to isolate rostro-caudal differences while controlling for sub-regional variability. Mapping these responses across the longitudinal axis of the PL confirmed a functional gradient that closely mirrored the reverse mPFC to BLA pathway, with rostral BLA inputs driving significantly larger EPSCs in the rostral PL, whereas caudal BLA inputs preferentially targeted the caudal PL (**Fig. 4f, i**).

These electrophysiological results demonstrate that the structural organisation of reciprocal prefrontal–amygdala loops is mirrored by distinct functional synaptic weights across the rostro-caudal axis. Given this topographical arrangement with strong reciprocal connectivity linking the rostral BLA and the rostral mPFC, we next tested its potential role in supporting fear memory formation.

### Post-learning activation of reciprocal BLA-mPFC connections drives the consolidated engram

Given the distinct topography of prefrontal–amygdala connectivity, we next investigated whether the post-consolidation activity shift in L2/3 of the mPFC reflects the selective engagement of long-range projection neurons that provide input to the BLA. Because the mPFC regulates fear memory retrieval via efferent projections to the BLA^22^, and these amygdala-projecting neurons are predominantly localised within superficial layers^37,38^, we reasoned that the elevated L2/3 activity during retrieval could represent a post-learning recruitment of this downstream pathway.

To test if fear memory consolidation coincides with enhanced activation of these downstream projections, we used the RAM system to drive expression of ChR2 in mPFC ensembles engaged during different stages of memory formation (**Fig. 5a-c**). To comprehensively assess the overall prefrontal input to the BLA across all conditions - and not to overlook localised shifts that might occur between different stages of learning - we targeted the entire mPFC ensemble without isolating rostral or caudal subregions (see Methods). We then obtained whole-cell patch-clamp recordings from principal neurons across the entire BLA and delivered blue light pulses to drive axon terminals of prefrontal cells captured during home cage, tone-only, fear learning, or fear retrieval states (**Fig. 5d**). The functional innervation rate of active cells with postsynaptic BLA neurons was highest when stimulating terminals from the fear retrieval ensemble (**Fig. 5e**). Furthermore, optogenetic stimulation of fear retrieval terminals evoked significantly larger EPSCs in BLA neurons compared to fear learning, home cage, or tone-only ensembles (**Fig. 5f**). Although the initial learning stage led to an increase in projection engagement relative to control conditions, the substantial increase observed following retrieval demonstrates that following memory consolidation there is a significant enhancement of mPFC drive of the BLA. In parallel, local microcircuit recordings in the mPFC revealed that the fear retrieval ensemble drives a marked increase in local inhibitory postsynaptic currents (IPSCs) and a shift toward dominant local inhibition onto neighbouring non-ensemble neurons (Extended Data Fig. 8). Overall, these findings show that the post-learning activation of superficial cells in the prefrontal cortex is associated with an enhanced engagement of long-range projections to the BLA.

**Figure 5.**
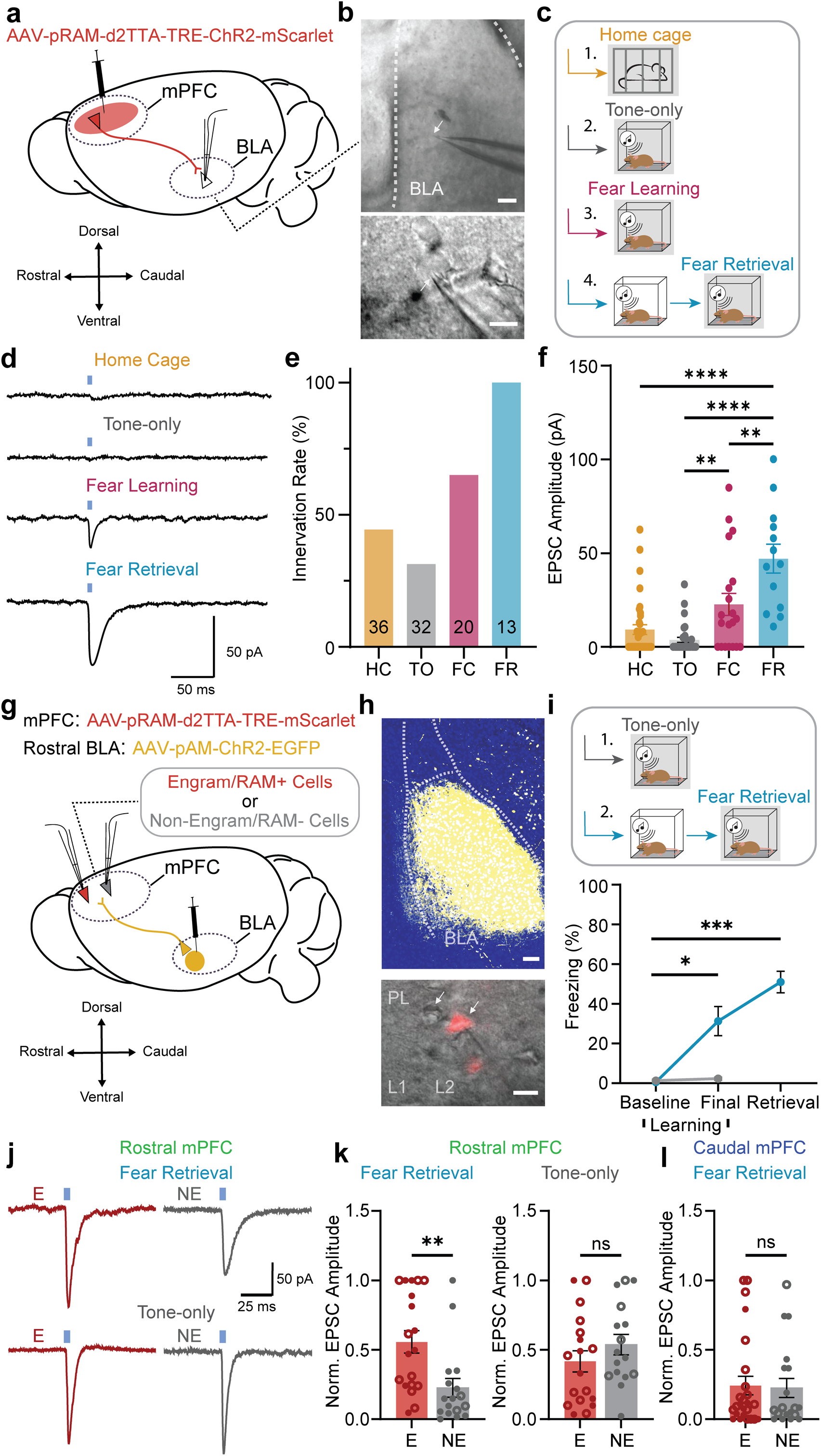
Post-learning engagement of reciprocal prefrontal–amygdala connectivity. **a.** Experimental strategy for activity-dependent terminal mapping. The RAM system was injected into the mPFC to drive ChR2 and mScarlet expression in cells active during specific behavioural windows. At least one-week post-tagging, acute brain slices were taken and whole-cell recordings were made from principal neurons in the BLA during optical reactivation of prefrontal axon terminals. **b.** Top: representative low-magnification DIC image of a recording site within the BLA; scale bar = 100. Bottom: high-magnification DIC image of a whole-cell patch-clamp recording from a principal BLA neuron; scale bar = 25 µm. **c.** The RAM system was used to capture active neuronal ensembles in the mPFC under four different behavioural conditions: home cage (HC, yellow, n = 4), tone-only (TO, grey, n = 4), fear learning (FC, purple, n = 5) and fear retrieval (FR, blue, n = 5). **d.** Representative light-evoked responses from BLA neurons when mPFC active ensemble axon terminals were optically stimulated across all behavioural conditions. Blue bars denote delivery of 5 ms, 470 nm light pulse. All recordings were performed at –60 mV. **e.** Functional innervation rate across groups, showing that optical reactivation of fear retrieval ensemble terminals recruits the highest percentage of BLA neurons. **f.** Evoked EPSC amplitudes recorded from BLA neurons across groups. The fear retrieval ensemble (n = 13 neurons) drove significantly larger EPSC amplitudes compared to fear learning (P < 0.01, n = 20), home cage (P < 0.0001, n = 36), and tone-only groups (P < 0.0001, n = 32; one-way ANOVA with Tukey’s test). The fear learning ensemble also drove larger responses than home cage and tone-only controls (P < 0.01). **g.** Experimental strategy for mapping rostral BLA inputs onto mPFC active ensembles. ChR2 (yellow) was injected into the rostral BLA, and RAM-mScarlet was injected into the mPFC. Whole-cell recordings were taken from active (RAM+; red) and neighbouring non-active (RAM–; grey) L2/3 neurons within the rostral (green) and caudal (blue) mPFC. **h.** Top: representative confocal image of ChR2 expression within the rostral BLA; scale bar = 100 µm. Bottom: high-magnification DIC image showing a patched RAM+ neuron in layer 2 of the PL adjacent to a subsequently recorded non-labelled RAM– neuron; scale bar = 25 µm. **i.** Top: experimental timeline for tagging of mPFC active ensembles during tone-only (n = 6 mice) and fear retrieval conditions (n = 6 mice). Bottom: freezing behaviour confirming significant conditioning (P < 0.05) and retrieval freezing (P < 0.001, two-way ANOVA with Šídák’s test) in the retrieval group, with no freezing increases in tone-only controls (P > 0.05). **j.** Representative EPSC traces recorded from active (RAM+; E) and non-active (RAM–; NE) neurons in the rostral mPFC during stimulation of rostral BLA inputs across fear retrieval (top) and tone-only conditions (bottom). **k.** Normalised EPSC amplitudes from active (E) and non-active (NE) neurons in the rostral mPFC. Open circles denote putative inhibitory interneurons; closed circles denote putative excitatory neurons. In fear retrieval animals, active neurons (n = 15) received significantly larger inputs from the rostral BLA compared to non-active neurons (n = 12; P < 0.01, unpaired t-test). No differences were observed in tone-only controls (n = 18, n = 16; P > 0.05). **l.** Normalised EPSC amplitudes recorded from active (n = 25) and non-active (n = 19) neurons within the caudal mPFC during stimulation of rostral BLA inputs, showing no significant differences (P > 0.05, unpaired t-test). All data are presented as mean ± SEM.

Next, we tested whether the prefrontal retrieval ensemble is driven by incoming inputs from the amygdala. Because the largest increase in activity was observed within the rostral PL, and this region receives most dominant input from the rostral BLA, we specifically targeted the rostral BLA to rostral mPFC circuit. We virally delivered ChR2 into neurons in the rostral BLA and the RAM system into the mPFC, tagging prefrontal cells active during either fear retrieval or a tone-only control session with mScarlet (**Fig. 5g-i**). Whole-cell patch-clamp recordings were then obtained from active (RAM+) and neighbouring non-active (RAM–) L2/3 principal neurons within the rostral or caudal PL (**Fig. 5j**).

In the rostral PL of fear retrieval animals, optogenetic stimulation of rostral BLA input evoked larger EPSCs in active cells than in adjacent non-active cells (**Fig. 5k**). This selective prioritisation in the rostral mPFC was absent in the tone-only control group, suggesting that following the formation of fear memory, the rostral BLA drive of PL retrieval ensemble neurons is enhanced (**Fig. 5k**). Crucially, this functional selectivity was absent within the caudal PL of fear retrieval animals, where active and non-active neurons received equivalent synaptic drive from the rostral BLA (**Fig. 5l**). Notably, mPFC L2/3 interneurons receiving strong BLA input (**Fig. 5k**, open circles) were biased toward active ensemble incorporation during fear retrieval relative to tone-only controls.

Collectively, these physiological profiles reveal that the post-consolidation ensemble is characterised by the coordinated engagement of bidirectional connectivity between the mPFC and BLA. The post-learning increase in superficial prefrontal activity is associated with an enhanced functional efferent drive to the amygdala. Concurrently, the rostral PL ensemble receives prioritised synaptic input from the rostral BLA, positioning this amygdalar subdivision as a key driver of the post-consolidation increase in prefrontal activity. Together, these physiological data demonstrate that the post-learning activation of reciprocal BLA – mPFC connections is a hallmark feature of the consolidated fear engram.

### Reciprocal connectivity between the rostral mPFC and rostral BLA supports the consolidated fear engram

Although the mPFC exhibits distinct anatomical gradients along its rostro-caudal axis, how this structural connectome shapes the functional organisation of the consolidated memory engram remains unknown. Specifically, it is not known whether the consolidated fear memory engages BLA-mPFC networks uniformly across this spatial axis, or if it preferentially engages topographical pathways. To evaluate if this rostro-caudal topography of reciprocal prefrontal-amygdala connectivity has functional relevance, we systematically mapped the efferent and afferent projections of the consolidated prefrontal fear engram. Using the RAM system to label the fear retrieval ensemble across the entire mPFC, we mapped the distribution of anterograde terminal fibres within the BLA (**Fig. 6a, b**). Although active cells were distributed across the entire mPFC (**Fig. 6c**), mapping their terminal fibres within the amygdala revealed a highly distinct pattern. The axonal projection distribution of the fear retrieval ensemble closely matched the restricted anatomical layout of the rostral mPFC, terminating preferentially within the rostral BLA (**Fig. 6b, d**). While our primary focus was the amygdala, comprehensive whole-brain mapping revealed that the fear retrieval engram also heavily innervates additional cortical and subcortical structures, including the agranular insular cortex (AI), claustrum (CLA), striatum (CP), and retrosplenial cortex (RSP) (Extended Data Fig. 9a-c). Thus, the output profile of the entire mPFC active ensemble closely matched the input distribution of the rostral mPFC to the BLA, suggesting that the rostral mPFC to rostral BLA pathway is preferentially engaged during fear memory retrieval.

**Figure 6.**
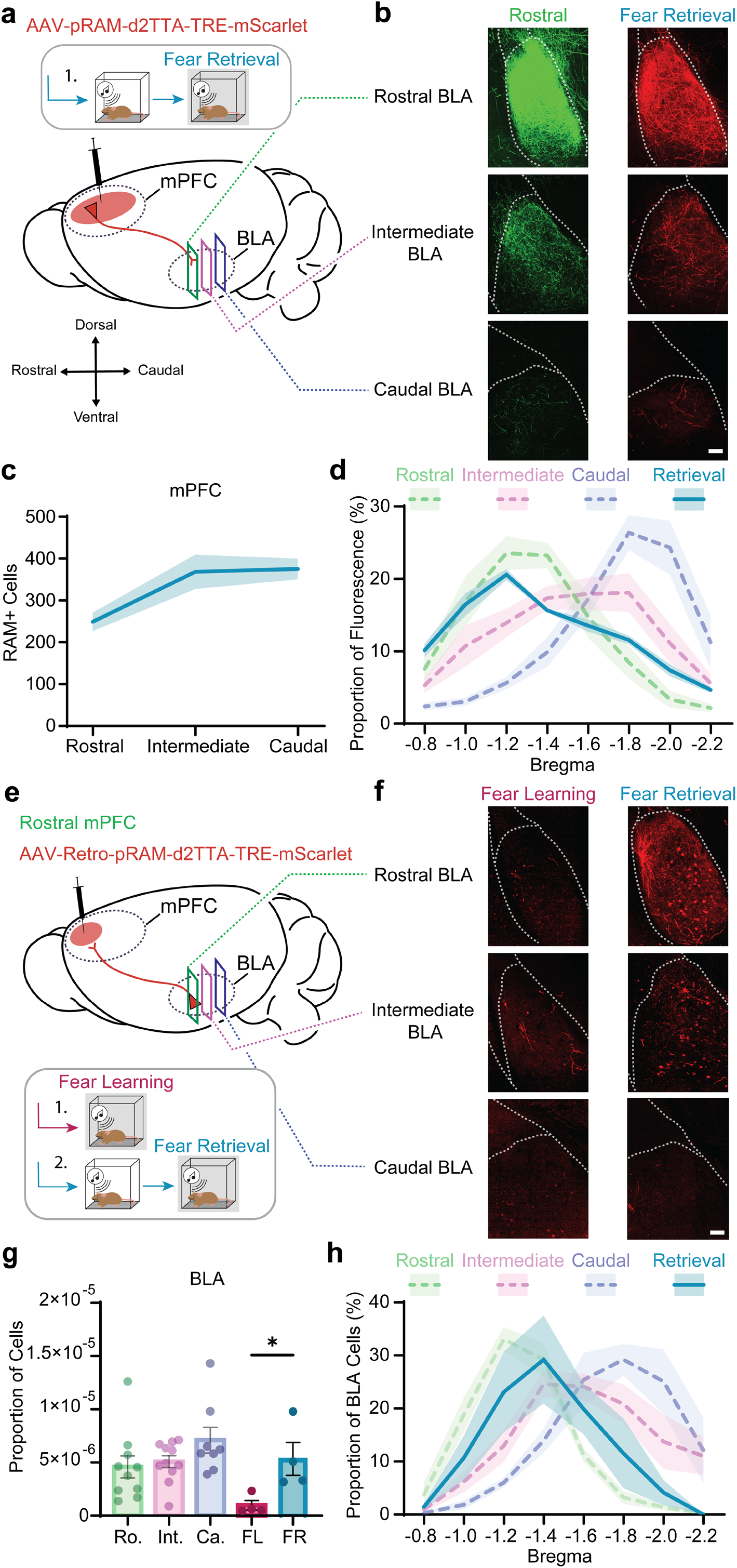
The consolidated fear ensemble is supported by a distinct reciprocal circuit between the rostral mPFC and rostral BLA. **a.** Experimental strategy for mapping the efferent output of the consolidated prefrontal fear ensemble. The RAM system was injected into the mPFC to express mScarlet in neurons active during fear retrieval. **b.** Left: baseline anatomical distribution of rostral mPFC efferent projections in the BLA (reproduced from Fig. 3 for comparison). Right: representative confocal images showing the axonal terminal fibres of the fear retrieval active ensemble within the rostral, intermediate, and caudal BLA. Scale bar = 100 µm. **c.** Mean number of active RAM+ cells across the rostral, intermediate, and caudal mPFC segments in animals included for efferent fibre analysis (n = 7). There were no significant differences in the total number of RAM+ cells across the three zones of the mPFC (P < 0.05, one-way ANOVA with Tukey’s test). **d.** Proportional distribution profile of RAM-specific terminal fluorescence across the rostro-caudal axis of the BLA (n = 7 mice; blue line) overlayed on baseline anatomical tracings from Fig. 3 (dotted lines). **e.** Experimental strategy for activity-dependent retrograde input mapping. Retro-RAM was virally delivered into the rostral mPFC to express mScarlet in active afferent projection neurons during either fear learning (n = 4) or fear retrieval (n = 4). **f.** Representative confocal images of retrogradely labelled active (RAM+) cells within the rostral, intermediate, and caudal BLA targeting the rostral mPFC during fear learning (left) and fear retrieval (right). Scale bar = 100 µm. **g.** Total area-normalised proportion of afferent cells projecting to the mPFC localised within the BLA. Fear retrieval produced a significant increase in the total number of active neurons projecting to the rostral mPFC, compared to fear learning (P < 0.05, unpaired t-test). Plotted alongside baseline anatomical proportions of cells projecting to the rostral (Ro), intermediate (Int) and caudal (Ca) mPFC (reproduced from Fig. 3 for comparison). **h.** Proportional distribution profile of active neurons projecting to the rostral mPFC across the rostro-caudal axis of the BLA during fear retrieval (n = 4 mice; blue line) overlayed on baseline anatomical tracings from Fig. 3 (dotted lines). All data are presented as mean ± SEM.

Next, we sought to determine whether the post-learning increase in rostral mPFC activity is associated with an enhanced functional input from the rostral BLA. To achieve this, we packaged the RAM system into a retrograde viral vector (retro-RAM) and delivered it into the rostral mPFC to label active projection neurons during fear learning or fear retrieval (see Methods; **Fig. 6e, f**). Quantifying labelled neurons across the BLA that project to the rostral mPFC revealed a significant increase in the number of active BLA projection neurons during fear retrieval compared to initial learning (**Fig. 6g**). Spatial mapping confirmed that these active projection neurons were concentrated within the rostral BLA, overlapping closely with our baseline anatomical inputs to the rostral mPFC (**Fig. 6h**). Beyond the amygdala, brain-wide afferent profiling revealed broader shifts in prefrontal input between learning and memory retrieval, marked by differences in active projection populations from areas such as the AI and anterior cingulate cortex (AC) (Extended Data Fig. 9d-f). In contrast, parallel retro-RAM injections into the caudal mPFC revealed significantly lower active BLA projection neurons during fear retrieval (Extended Data Fig. 9g-j). These findings demonstrate that fear memory consolidation is accompanied by an increase in active projection neurons principally within the rostral BLA to rostral mPFC circuit during retrieval.

Ultimately, by mapping functional distributions back onto the underlying structural framework, these results demonstrate that the post-consolidation fear memory ensemble is characterised by the selective engagement of a bidirectional circuit between the rostral mPFC and the rostral BLA. The downstream efferent profile of the prefrontal ensemble aligns closely with that of the rostral mPFC, while its post-learning activation is mirrored upstream by an increased active input from the rostral BLA.

### Rostral mPFC ensembles are necessary and sufficient for fear memory expression and require input from the rostral BLA

Classical frameworks suggest that memory retrieval is underpinned by the reactivation of specific neuronal networks established during learning and consolidation^7,44^. Given the pronounced post-learning engagement of reciprocal rostral mPFC – rostral BLA synaptic connections, we reasoned that this localised prefrontal ensemble actively drives fear memory expression. However, whether this specific network is causally necessary or sufficient for memory retrieval remains to be established.

To evaluate whether this prefrontal ensemble is required for memory expression, we utilised the RAM system to express the inhibitory opsin archaerhodopsin (ArchT) or a control fluorophore in rostral mPFC cells active during an initial retrieval session (see Methods; **Fig. 7a-c** and Extended Data Fig. 10). During a subsequent retrieval test in the conditioned context, direct optogenetic suppression of this retrieval-tagged ensemble via continuous green light delivery during CS presentations, elicited a significant reduction in freezing compared to controls (**Fig. 7d**). These data demonstrate that the activation of this specific post-consolidation network is necessary for the physiological retrieval of fear memory.

**Figure 7.**
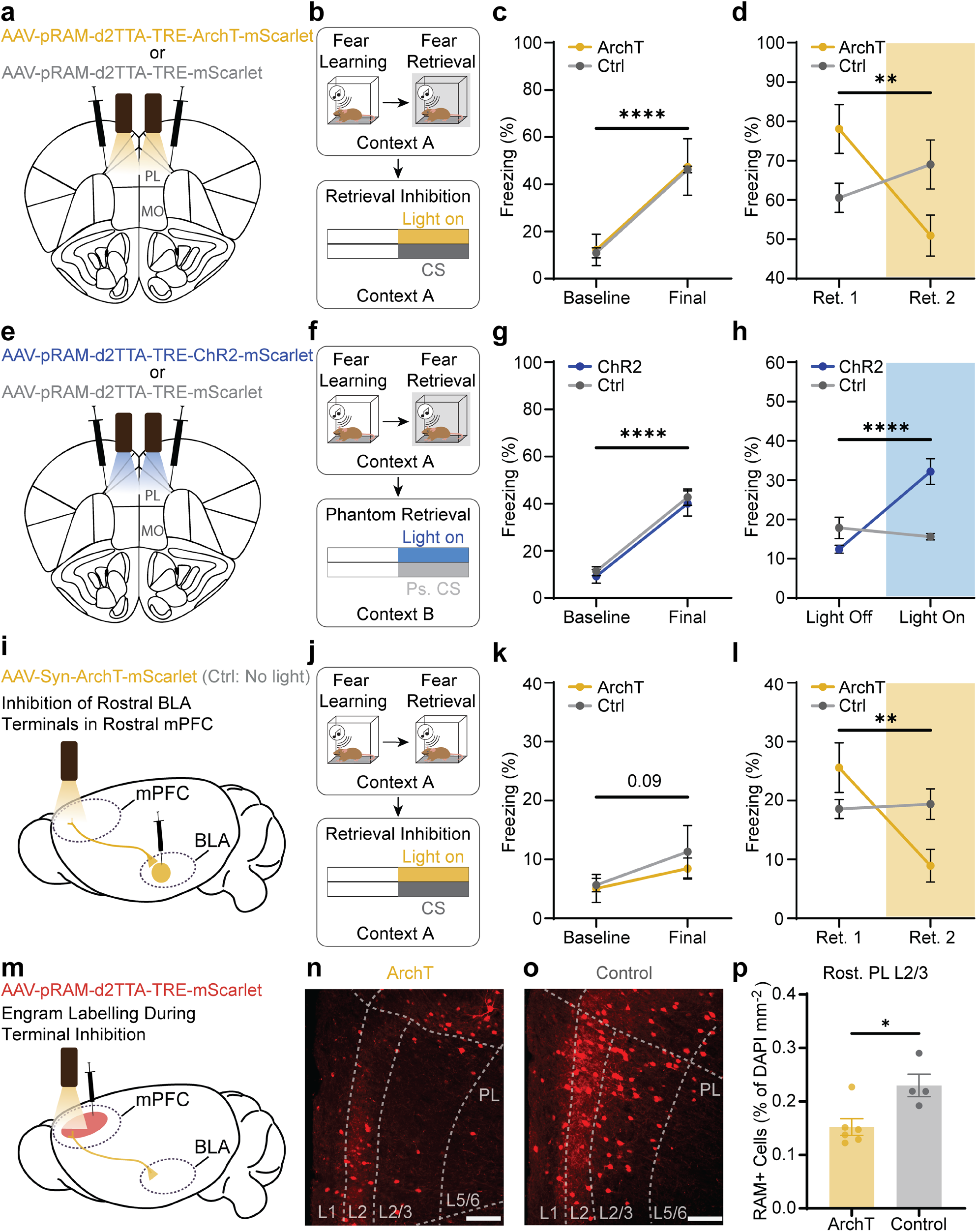
Consolidated prefrontal active ensembles are necessary and sufficient for fear memory expression. **a.** Strategy for optogenetic necessity testing of the mPFC fear retrieval ensemble. The RAM system was virally delivered into the mPFC to express ArchT and mScarlet (active group; n = 4) or mScarlet alone (control group; n = 7) in cells active during an initial retrieval test (Ret. 1). Bilateral optical fibres were implanted over the rostral PL (+2.4 mm Bregma). **b.** Experimental schematic. Four days after tagging cells during Ret. 1 in Context A, animals were returned to Context A for a second retrieval test (Ret. 2), during which CS presentations were coupled with continuous green light illumination (10 s, 560 nm). **c.** Freezing behaviour during conditioning, showing equivalent fear acquisition across active and control groups (P < 0.0001, two-way ANOVA). **d.** Freezing behaviour during Ret. 1 (baseline tag) and Ret. 2 (expression). Direct optogenetic suppression of the somatic retrieval ensemble drove a significant reduction in freezing behaviour during Ret. 2 compared to controls (P < 0.01, two-way ANOVA with Šídák’s test). **e.** Strategy for optogenetic sufficiency testing of the mPFC fear retrieval ensemble. The RAM system was injected bilaterally into the mPFC to express ChR2 and mScarlet (active group; n = 7) or mScarlet alone (control group; n = 7) in neurons active during fear retrieval. Bilateral optical fibres were implanted over the rostral PL (+2.4 mm Bregma). **f.** Experimental schematic. Two days after learning, mPFC ensembles were tagged during a retrieval test in Context A. Four days later, animals were exposed to a phantom retrieval test in a novel context (Context B), during which natural tones were replaced with blue light stimulation pulses (15 ms, 30 Hz, 10 s, 473 nm) with the same time structure as the pseudo-CS (Ps. CS). **g.** Freezing behaviour during conditioning, showing equivalent and robust fear acquisition across groups (P < 0.0001, two-way ANOVA). **h.** Freezing behaviour during the phantom retrieval test. Optogenetic reactivation of the retrieval-tagged prefrontal ensemble drove a significant increase in freezing behaviour in the active group during light on epochs compared to the pre-stimulation baseline period (Light Off; P < 0.0001, two-way ANOVA with Šídák’s test). No significant differences in freezing behaviour were observed across light off and on epochs in the control group (P > 0.05). **i.** Strategy for terminal pathway necessity testing for driving the mPFC fear engram ensemble. ArchT was virally delivered bilaterally into the rostral BLA, and bilateral optical fibres were implanted over terminal fields within the rostral/intermediate PL (+2.4 mm Bregma). To control for viral delivery variables, both the active group (light delivered; n = 6) and the control group (no light delivered; n = 5) received the same ArchT construct. **j.** Experimental schematic. Following fear conditioning, animals underwent a baseline retrieval test (Ret. 1) in Context A in the absence of light, followed two days later by a second retrieval test (Ret. 2) in Context A, during which CS presentations were coupled with terminal optogenetic inhibition via continuous green light (10 s, 560 nm). **k.** Freezing behaviour during fear conditioning, showing increased and equivalent fear acquisition curves across active and control groups (P = 0.09, two-way ANOVA). **l.** Freezing behaviour during baseline retrieval (Ret. 1) and terminal inhibition (Ret. 2). Direct optogenetic suppression of incoming rostral BLA terminals within the rostral mPFC during CS presentation produced a significant reduction in freezing behaviour compared to the control condition (P < 0.01, two-way ANOVA with Šídák’s test). All data are presented as mean ± SEM. **m.** Experimental strategy for activity dependent labelling of active cells in the mPFC during inhibition of rostral BLA terminals in the rostral mPFC (Ret. 2). **n.** Example confocal image of RAM labelled neurons in the rostral PL across L2/3 and L5/6 during the retrieval of fear memory in the ArchT group. Scale bar = 100 μm. **o.** Example confocal image of RAM labelled neurons in the rostral PL across L2/3 and L5/6 during the retrieval of fear memory in the control group. Scale bar = 100 μm. **p.** Mean number of active RAM-labelled cells in L2/3 of the rostral PL (% of DAPI/mm^2^). Inhibition of rostral BLA terminals in the rostral mPFC was associated with a significant reduction in active cells in L2/3 compared to the control condition (P < 0.05, unpaired t test).

Having established necessity, we next evaluated whether this rostral mPFC ensemble is sufficient to drive fear memory expression. We used the RAM system to express the excitatory opsin ChR2, or a control fluorophore, in prefrontal cells active during an initial fear retrieval test (see Methods; **Fig. 7e-g** and Extended Data Fig. 10). Four days later, animals were placed into a novel context and exposed to a phantom retrieval test, where natural tones were replaced with blue light stimulation delivered through bilateral optic fibres implanted over the rostral PL (**Fig. 7e, f**). Optogenetic reactivation of the retrieval-tagged prefrontal ensemble evoked a significant increase in freezing, demonstrating that the artificial reactivation of this prefrontal network is sufficient to drive fear memory expression (**Fig. 7h**).

Finally, we evaluated the necessity of the upstream BLA input driving this prefrontal representation. We virally expressed ArchT bilaterally in the rostral BLA and implanted bilateral optical fibres over terminal fields within the rostral mPFC (see Methods; **Fig. 7i-k**). During testing, optogenetic suppression of rostral BLA terminals in the rostral mPFC during CS presentations produced a significant reduction in freezing compared to the control condition (**Fig. 7l**). Crucially, post-hoc quantification revealed that this real-time terminal inhibition also caused a reduction in the number of active L2/3 cells labelled by the RAM system within the rostral PL during retrieval (**Fig. 7m-p**).

In summary, these data demonstrate that the ensemble within the rostral mPFC is necessary and sufficient for fear memory expression. Moreover, these findings establish that monosynaptic input from the rostral BLA to the rostral mPFC is essential during retrieval, indicating that acute suppression of this distinct pathway is sufficient to impair the retrieval of consolidated fear memory.

## Discussion

Our findings identify a topographically organised, reciprocal circuit linking the rostral BLA and rostral mPFC that drives the expression of consolidated fear memory. By tracking active networks across distinct stages of memory processing, we show that the retrieval of a fear memory does not simply reactivate the same cells engaged during learning; rather, there is a post-learning engagement of L2/3 cells concentrated along a spatial gradient in the rostral PL cortex (**Figure 1, 2**). This prefrontal ensemble is both necessary and sufficient to drive fear memory retrieval (**Figure 7**), and its maintenance relies on a functional gradient biased toward input from the rostral BLA (**Figure 5, 6**).

These data provide a structural and functional framework that contextualises historical discrepancies regarding prefrontal contributions to fear conditioning. Prior studies utilising single-unit recordings in the PL reported contradictory findings, describing either broad decreases or marked increases in firing rates following fear conditioning^24,45^. Our anatomical and functional mapping demonstrates that these divergent observations are likely a consequence of treating the PL as structurally and functionally homogeneous along its longitudinal axis. By systematically mapping connectivity across the rostro-caudal axis, we show that prefrontal–amygdala communication is organised in topographically distributed loops (**Figure 3, 4**). Within this architecture, superficial layers of the rostral PL maintain dense reciprocal connectivity with the rostral BLA. Although prior work has noted target preferences between mPFC subregions^41,46^, our results reveal a broader circuit organisational principle where the rostral PL receives dense innervation from the rostral BLA. This structural layout translates directly into a functional gradient during memory retrieval, where neuronal engagement is heavily concentrated within the rostral PL and progressively tapers off toward more caudal prefrontal regions.

This topographical distribution may represent a wider rule for prefrontal-amygdala interactions, with different information processed in parallel, but distinct circuits. Under this framework, core threat information is processed and stored within a rostro-dorsal module linking the rostral PL to the rostral BLA, whereas safety or extinction memories could engage parallel connections linking the caudo-ventral mPFC to the caudal BLA. The functional organisation of these loops may be guided by an underlying segregation of valence processing within the amygdala, where negative-valence, shock-responsive neurons dominate the rostral BLA and positive-valence, extinction-responsive neurons populate the caudal BLA^47,48^. These distinct subcortical populations have been shown to mutually antagonise each other through local feed-forward inhibitory circuits^47,49^. Consequently, extinction learning selectively recruits positive-valence neurons in the caudal BLA, which could suppress rostral fear networks and engage projections to the caudo-ventral prefrontal cortex (i.e. infralimbic cortex)^34,48^, with the spatial architecture of these loops governing valence assignment and subsequent memory expression.

The laminar shift toward L2/3 prefrontal activity during retrieval also has implications for models of cortical computation, and systems consolidation. Neurons in L2/3 of the mPFC are primary recipients of long-range associative afferents and act as central hubs for local intracortical processing^36,37,40,50^. The selective enrichment of functional synaptic input from the rostral BLA onto retrieval-tagged L2/3 prefrontal neurons indicates that long-term memory stabilisation involves the engagement of subcortical-to-cortical associative connections (**Figure 5**). Furthermore, as optogenetic stimulation of these prefrontal retrieval cells drives significantly larger synaptic responses in the BLA than stimulation of learning ensembles, our data indicate that consolidation may establish a highly efficient bidirectional feedback loop^18,51^. This prefrontal engram incorporates a significant population of L2/3 GABAergic interneurons that receive robust, direct excitatory drive from the BLA and execute strong local inhibition onto surrounding non-engram principal cells (**Figure 5** and Extended Data Fig. 2, 8). BLA inputs have been shown to drive feed-forward inhibition through somatostatin-expressing (SOM+) interneurons that synapse onto L2/3 mPFC neurons projecting back to the amygdala^37,40^, a circuit layout refined by learning-induced plasticity at BLA-to-SOM synapses^52^. Thus, the post-learning engagement of reciprocal connectivity between the mPFC and BLA could be supported by local microcircuitry that coordinates population activity between regions.

Importantly, our finding that the prefrontal network engaged during retrieval of an auditory fear memory is distinct from the ensemble engaged during learning challenges traditional interpretations of the engram. The low cellular overlap typically observed between learning and retrieval ensembles^13,14,16,17^ is not necessarily an experimental artifact. Instead, it may reflect a fundamental feature of systems memory consolidation, during which the physical trace is actively transformed and distributed across specialised circuits^1,3,53^. This dynamic reorganisation aligns with our data reflecting a temporal pattern of circuit engagement where initial learning is a BLA-dependent process, whereas subsequent retrieval requires the progressive recruitment of cortical nodes. Crucially, initial fear learning requires temporally aligned activation of conditioned and unconditioned inputs that occurs primarily within the lateral amygdala (LA)^54–57^, which has very few direct connections with the mPFC^27^. Therefore, maturation of the memory trace may require a shift from these dorsal encoding sites to basal regions of the BLA, which possess the dense reciprocal connections required for subsequent expression, a transition likely mediated by internal intra-amygdalar networks^58,59^. In combination with local microcircuitry, these long-range bidirectional networks represent an ideal substrate for offline processing, structurally optimised to sustain persistent representations and entrain population activity over extended intervals^60,61^. This reciprocal circuit structure could drive synchronised, offline replay events across the rostral BLA – mPFC loop during post-learning sleep windows, providing the necessary recurrent interaction to drive synaptic plasticity during sleep^4,62–65^ and serving as a core circuit mechanism underpinning systems memory consolidation. Identifying the exact molecular and synaptic mechanisms that alter these parallel pathways during the consolidation window will be a critical step toward understanding long-term memory stabilisation and storage.

## Methods

### Animals

All experiments were approved by the Animal Ethics Committee of The University of Queensland in accordance with the Australian Code of Practice for the care and use of animals for scientific purposes. Adult male and female C57BL/6J mice (6–10 weeks of age) were housed under a standard 12-hour light/dark cycle with ad libitum access to food and water. For engram tagging experiments, animals were maintained on a diet supplemented with doxycycline (Dox; 625mg/kg or 40mg/kg) for a minimum of 2 days prior to surgery and throughout the behavioural paradigm, except during specified tagging windows. For experiments investigating the overlap between RAM, c-Fos and inhibitory neuron populations, GAD67-EGFP were used to endogenously label inhibitory neurons.

### Surgical procedures

Two days prior to surgery, animals were placed onto high doxycycline feed (625 mg/kg), to prevent activation of the RAM system during the period immediately after surgery. On the day of surgery, animals were anaesthetised with intraperitoneal injection of ketamine (100 mg/kg)/xylazine (10 mg/kg) and mounted in a stereotactic frame. The skull was exposed through a midline incision and small holes drilled over the desired injection site. Then, 0.4-0.6 µL of AAV-pRAM-d2TTA-TRE-mScarlet (4.36×10^13^ vg/mL) was injected unilaterally into the intermediate mPFC (A/P: +2.2 mm, M/L: ±0.3 mm, D/V:-1.1 mm) at a rate of 0.1–0.2 μL/min using 30-gauge needles coupled to a 5-μL Hamilton syringe, to allow spread of virus across all sub-regions and the rostro-caudal axis. The needle was slowly removed 5 min after injection was completed and the incision was then disinfected and closed using Vetbond tissue adhesive. Animals were administered Metacam (4 μL/10 g) diluted in 1 mL of saline injected subcutaneously. A minimum of three weeks was maintained between surgery and the commencement of experimental procedures to allow for recovery and viral expression. For the blocking of NMDA receptors during fear learning, the injection of the RAM system in the mPFC was supplemented with the mounting of bilateral infusion cannulas over the rostral BLA (A/P:-1.1 mm, M/L: ±3.0 mm, D/V:-3.7 mm).

For the investigations of long-range and local circuitry of mPFC active populations, the RAM system was used to drive the expression of the excitatory opsin, ChR2, in cells that were active during the behavioural window. Therefore, 0.4-0.6 µL of AAV-pRAM-d2TTA-TRE-ChR2-mScarlet (4.63×1013 vg/mL) was injected unilaterally into the intermediate mPFC. To trace the efferent projections from the rostral, intermediate and caudal mPFC, we injected 0.05-0.1 µL of AAV-pAM-EGFP (4.91×10^12^ vg/mL) into the rostral mPFC (A/P: +2.6 mm, M/L: ±0.3 mm, D/V:-0.9 mm), 0.05-0.1 µL of AAV-pAM-tdTomato (2.81×10^13^ vg/mL) into the intermediate mPFC (A/P: +2.2 mm, M/L: ±0.3 mm, D/V:-1.1 mm), and 0.05-0.1 µL of AAV-pAM-mTagBFP2 (6.28×10^12^ vg/mL) into the caudal mPFC (A/P: +1.7 mm, M/L: ±0.3 mm, D/V:-1.2 mm). To trace the afferent projections to the rostral, intermediate and caudal mPFC, we injected 0.05-0.1 µL of AAV-Retro-pAM-EGFP (2.14×10^12^ vg/mL) into the rostral mPFC (A/P: +2.6 mm, M/L: ±0.3 mm, D/V:-0.9 mm), 0.05-0.1 µL of AAV-Retro-pAM-pemiRFP670-N1 (6.99×10^12^ vg/mL) into the intermediate mPFC (A/P: +2.2 mm, M/L: ±0.3 mm, D/V:-1.1 mm), and 0.05-0.1 µL of AAV-Retro-pAM-mTagBFP2 (8.25×10^12^ vg/mL) into the caudal mPFC (A/P: +1.7 mm, M/L: ±0.3 mm, D/V:-1.2 mm). These injection sites are the mid-point of each anatomical mPFC location used to group active cells in various stages of fear learning (Rostral: +2.8 - 2.4 mm Bregma; Intermediate: +2.3 - 2.0 mm Bregma; Caudal: +1.9 - 1.5 mm Bregma). Rather than targeting a single sub-region of the mPFC, each retrograde virus was targeted at the mid-point on the dorso-ventral axis for that rostro-caudal location. However, only those injection sites that had expression that included the PL were used for analysis, to enable consistency of a single sub-region across all injection sites. To account for the potential bias of the virus arrangement, a small portion of the anterograde (n = 2 of 7) and retrograde (n = 4 of 12) cohorts had the location of each virus randomised in their order.

For mapping convergent inputs to the BLA from the rostral and caudal mPFC using iDOTS, we injected AAV-pAM-ChR2-mScarlet (5.67×10^12^ vg/mL) into the rostral mPFC (A/P: +2.6 mm, M/L: ±0.3 mm, D/V:-0.9 mm) and AAV-stChrimson-PdCO-YFP into the caudal mPFC (A/P: +1.7 mm, M/L: ±0.3 mm, D/V:-1.2 mm). For electrophysiological investigation of synaptic inputs from the BLA to mPFC, 0.1-0.2 µL of AAV-pAM-ChR2-EGFP (7.90×10^12^ vg/mL) was injected into the rostral BLA (A/P:-1.1 mm, M/L: ±3.0 mm, D/V:-3.7 mm) or caudal BLA (A/P:-1.7 mm, M/L: ±3.1 mm, D/V:-3.8 mm). For investigations of synaptic input from the BLA onto active and non-active mPFC neurons, 0.1-0.2 µL of AAV-pAM-ChR2-EGFP was injected into the rostral BLA (same coordinates) and 0.4-0.6 µL of AAV-pRAM-d2TTA-TRE-mScarlet (4.36×10^13^ vg/mL) was injected into the intermediate mPFC (A/P: +2.2 mm, M/L: ±0.3 mm, D/V:-1.1 mm).

For the labelling of active cells projecting to the mPFC during fear learning and retrieval, 0.2-0.3 µL of AAV-Retro-pRAM-d2TTA-TRE-mScarlet (1.77×10^13^ vg/mL) was injected into the rostral mPFC (A/P: +2.6 mm, M/L: ±0.3 mm, D/V:-0.9 mm) or caudal mPFC (A/P: +1.8 mm, M/L: ±0.3 mm, D/V:-1.2 mm).

For optogenetic reactivation or inhibition of engram cells *in vivo*, 0.4-0.6 µL of either AAV-pRAM-d2TTA-TRE-ChR2-mScarlet (active ChR2 group), AAV-pRAM-d2TTA-TRE-ArchT-mScarlet (active inhibition group) or AAV-pRAM-d2TTA-TRE-mScarlet (4.36×10^13^ vg/mL) (control group) was injected bilaterally into the border between the rostral and intermediate mPFC zones (A/P: +2.4 mm, M/L: ±0.3 mm, D/V:-0.9 mm). Then, bilateral optic fibres (Doric lenses) were implanted in the PL 0.1 mm above the injection site and fixed with dental cement. The sides of the optic fibres were painted with dried green retrobeads to allow post hoc analysis of fibre location. The skin around the implant was then glued using VetBond to the hardened dental cement to prevent infection. For the optogenetic inhibition of rostral BLA terminals in the rostral mPFC, 0.1-0.2 µL of AAV-Syn-ArchT-mScarlet (9.59×10^11^ vg/mL) was injected bilaterally into the rostral BLA (A/P:-1.2 mm, M/L: ±3.0 mm, D/V:-3.7 mm), and bilateral optic fibres implanted in the rostral PL (A/P: +2.4 mm, M/L: ±0.3 mm, D/V:-0.9 mm). Due to the effect of expressing inhibitory viral constructs in the rostral BLA on fear learning, both the active and control group received the same viral construct, while only the active group received light stimulation to isolate the effect of inhibiting rostral BLA terminals in the rostral mPFC.

### Auditory fear conditioning and engram tagging

One week prior to the behaviour of interest, animals were placed on a low concentration doxycycline feed (40 mg/kg) to lower doxycycline levels in the system to allow for tagging during the behavioural window. For the tagging of fear learning cells with RAM, animals were taken off doxycycline feed 48 hours prior to the learning session. Auditory fear conditioning consisted of five 10 second tones (4 kHz), which co-terminated with a 0.5 sec, 0.5 mA foot-shock (Coulbourn Instruments). For tagging of fear retrieval cells, animals received fear conditioning while on doxycycline feed and were taken off this feed immediately after the learning session. Then, 48 hours later, animals were placed back into the learning environment and received two 10 second tones (4 kHz). For the tone-only control group, identical procedures were performed as for the fear conditioning group, but the foot shocks were omitted, and animals only received five tone presentations. For the home cage control group, animals were taken off doxycycline for a 48-hour period and left in the home cage. For the tagging of remote retrieval cells in the mPFC, the interim period between fear conditioning and retrieval was lengthened to seven days. Thus, these animals were taken off doxycycline feed five days after the learning session. All animals were immediately placed back onto a high concentration doxycycline feed (625 mg/kg). For differential conditioning experiments, animals received 3 consecutive days of conditioning with the presentation of five CS+ tones (4 kHz), each paired with a foot-shock (0.5 sec, 0.5 mA), as well as five CS-tones (10 kHz) which were not associated with a foot-shock. The presentation of the CS+ and CS-were randomised in their order. Two days after the final conditioning session, animals received a retrieval test of two presentations of the CS+ or CS-, while off doxycycline feed. For the blocking of NMDA receptors during fear learning, APV (50 mM) or saline was bilaterally infused into the BLA 30 minutes prior to fear conditioning.

### Optogenetic manipulations

For *in vivo* optogenetic activation, animals were connected to a laser diode module via a patch cord and rotary joint. For phantom memory retrieval, retrieval-tagged ChR2 animals were placed into a novel context (Context B; plexiglass floor, ambient lighting, cleaned with 1% acetic acid). Following a baseline period, animals received blue light stimulation pulses (15 ms pulses, 30 Hz, 10 s duration, 473 nm, ∼10–12 mW at fibre tip) delivered at timelines corresponding to the original CS presentations.

For *in vivo* optogenetic inhibition, green light (continuous, 10 s duration, 560 nm, ∼15 mW at fibre tip) was delivered bilaterally into the prefrontal cortex during CS presentations to activate ArchT, suppressing either retrieval-tagged somata or incoming axon terminals originating from the rostral BLA. In both cases, retrieval sessions were separated to allow for tagging with the RAM system. In the RAM-ArchT inhibition of fear retrieval cells in the mPFC, the initial retrieval session (Ret. 1) is used to tag the ensemble underpinning the natural expression of the memory. This is then followed by the inhibition retrieval session (Ret. 2) after allowing for opsin expression, whereby fear retrieval cells are inhibited during CS presentations. Similarly, for inhibition of rostral BLA terminals in the rostral mPFC, the first retrieval session (Ret. 1) is separated from the light stimulated retrieval session (Ret. 2) to allow for concurrent tagging of the mPFC fear retrieval ensemble in the mPFC while rostral BLA terminals are suppressed.

### Whole-cell patch-clamp electrophysiology

For all electrophysiological experiments, animals were anaesthetised with isoflurane (1mL in an enclosed container) and sacrificed through decapitation. Brains were quickly removed and placed into a cold cutting solution containing (in mM): NaCl 118, KCl 2.5, NaHCO3 25, glucose 10, MgCl2 4, CaCl2 0.5, and NaH2 PO4 1.2. Coronal slices (300 µm) were taken using a vibratome (Leica VT 1000S) and placed into oxygenated (95% O_2_/5% CO_2_) artificial cerebrospinal fluid (aCSF) containing (in mM): NaCl 118, KCl 2.5, NaHCO3 25, glucose 10, MgCl2 1.3, CaCl2 2.5, and NaH2 PO4 1.2. Slices were incubated at 32-34°C for 30-60 mins and then allowed to rest at room temperature for at least 30 mins before recordings were taken. Slices were transferred to a recording chamber perfused with a constant flow of oxygenated aCSF maintained at 32-34°C. Slices were visualised using an upright microscope (BX50WI, Olympus Optical, Tokyo, Japan) with a 5× NA 0.1 or 40× NA 0.8 objective and with infrared and differential interference contrast optics. Fluorescent neurons were visualised using an LED system (pE-2, CoolLED) and RFP filter sets (Olympus). Electrodes (3–7 MΩ) were filled with a pipette solution containing (mM): KMeSO4 135, NaCl 7, HEPES 10, Mg2 ATP 2, Na3GTP 0.3, and biocytin 8 (pH 7.3 with KOH, osmolarity ∼290 mOsm/kg). Signals were recorded using a patch clamp amplifier (Multiclamp 700B, Axon instruments). Signals were filtered at 10 kHz and digitised at 20 kHz (Instrutech, ITC-16). All data were acquired, stored, and analysed using Axograph X (Axograph, V 1.8.0). Recorded currents were low-pass filtered at 2 kHz and a minimum of 10 response sweeps were averaged for analysis. Firing properties were recorded in current-clamp mode through injection of a square current pulse increasing in 25 pA increments. IPSC and EPSC amplitudes were analysed as the peak amplitude in the period after the light pulse, while neuron type was determined through the action potential amplitude and spike width at half amplitude, at two times the action potential threshold (2T). Excitatory neurons were considered as those with action potential amplitudes >100 mV and a spike width >1 ms. Neurons with action potential amplitude and spike width below these thresholds were considered putative inhibitory interneurons.

For independent Dual-Opsin Terminal Stimulation (iDOTS) mapping of mPFC-to-BLA projections, whole-cell patch-clamp recordings were performed from postsynaptic neurons distributed across the longitudinal axis of the basolateral amygdala (BLA; Bregma –0.9 mm to –2.1 mm). Principal neurons were held at –60 mV in the voltage-clamp configuration to isolate evoked excitatory postsynaptic currents (EPSCs). To independently recruit converging inputs from the rostral medial prefrontal cortex (mPFC; expressing ChR2-mScarlet) and the caudal mPFC (expressing Chrimson-PdCO-YFP) onto the same postsynaptic cell without spectral crosstalk, an optical pulse sequence was delivered through the microscope objective. Caudal inputs were isolated by a 5 ms pulse of 640 nm light to activate Chrimson at caudal prefrontal terminals, and peak EPSC amplitudes were quantified within a locked post-stimulation window. To shunt subsequent transmitter release from these same caudal terminals, a 100 ms pulse of 405 nm light was applied shortly after to activate the inhibitory opsin PdCO. A second 5 ms pulse of 640 nm light was used to check the suppression of caudal mPFC terminals near the peak of the Chrimson excitation spectrum. A subsequent 5 ms pulse of 405 nm light was then delivered to selectively activate ChR2 at rostral prefrontal terminals during the ongoing period of caudal presynaptic inhibition. Finally, a long 3 second pulse of 560 nm was applied to deactivate PdCO (i.e. release the inhibitory control of caudal mPFC inputs) and reset the configuration for the subsequent sweep. The functional innervation rate was defined as the percentage of recorded neurons demonstrating responses driven exclusively by the rostral mPFC, caudal mPFC, convergent inputs from both subdivisions, or no detectable input. For functional mapping of the reverse BLA-to-mPFC pathway, whole-cell patch-clamp recordings were obtained at a holding potential of –60 mV from principal neurons across prefrontal sub-regions (prelimbic, PL; medial orbital, MO; cingulate, Cg; and infralimbic, IL cortices) and cortical layers (L2/3 and L5/6). To resolve the functional gradient along the longitudinal plane of the PL, recordings were systematically sampled across the anterior-posterior axis of L2/3 PL neurons (Bregma +2.6 mm to +1.7 mm). Monosynaptic EPSCs were evoked by optogenetic stimulation of ChR2-expressing axonal terminals originating from either the rostral or caudal BLA using a 5 ms pulse of 470 nm light. For all mapping configurations, peak EPSC amplitudes were extracted and analysed using Axograph X, and amplitudes were normalised within-animal prior to group-level topographical mapping and statistical evaluation to control for variability in viral transduction efficiency across cohorts.

For functional mapping of prefrontal efferent projections, whole-cell patch-clamp recordings were obtained from principal neurons within the BLA at a holding potential of –60 mV in the voltage-clamp configuration. To evaluate changes in functional connectivity across distinct stages of memory processing, prefrontal axon terminals originating from active ensembles tagged during specific behavioural windows (home cage, tone-only, fear learning, or fear retrieval) were photostimulated via blue light pulses (5 ms, 470 nm, 0.2 Hz) delivered through the objective. The functional innervation rate was defined as the percentage of recorded BLA neurons demonstrating detectable light-evoked monosynaptic EPSCs. For mapping the local connectivity of active ensembles in the mPFC, acute coronal slices were taken from the mPFC in the same animals as those used for efferent connections to the BLA. Whole-cell patch-clamp recordings were taken from non-labelled (RAM-) cells across the mPFC and EPSCs/IPSCs evoked using blue light pulses (5 ms, 470 nm, 0.2 Hz). EPSCs were evoked at a holding potential of-60 mV, while IPSCs were stimulated at-40 mV. The ratio of excitation and inhibition for each neuron recorded in the mPFC was calculated as the IPSC amplitude divided by the EPSC amplitude. Therefore, a non-labelled neuron that only received local inhibitory input from the active ensemble would have a value of 1, while a neuron that only received excitatory input would have a value of 0.

For assessing synaptic input from the rostral BLA onto active and non-active mPFC neurons, whole-cell recordings were obtained from RAM+ and RAM-neurons from the same slices, under consecutive matched conditions; that is, recording from one type of neuron was followed by the recording from a neuron of the other type within 50 µm of the first, within the same cortical layer (e.g. a RAM-neuron in L2, immediately followed by a RAM+ neuron in L2). As recordings from active and non-active neurons are not simultaneous in time, for statistical purposes these are considered unpaired recordings. Again, synaptic responses were stimulated using blue light pulses (5 ms, 470 nm) at-60 mV. Responses were grouped into rostral and caudal mPFC categories based on the position of the slice along the rostro-caudal axis. For this experiment, slices were assigned to the rostral mPFC when they were between approximately +2.8 and +2.3 Bregma, while the caudal mPFC was considered as slices between approximately +2.2 and +1.7 Bregma. This slightly wider range was implemented due to the inconsistencies of cutting acute slices of the mPFC, removing the intermediate mPFC as a separate category from analysis in this instance. Excitatory synaptic responses were normalised to the largest peak EPSC response for each individual animal, to remove the variance associated with differences in rostral BLA opsin expression across animals.

Two days after the optogenetic reactivation of mPFC engram cells, brain slices from behavioural animals expressing ChR2 in RAM neurons were prepared according to the protocol described above. Whole-cell recordings were obtained from RAM infected neurons immediately below the fibre tip, determined by the green fluorescence from the dried retrobeads, and ChR2 expression tested using 100 ms pulses of 470 nm light. In addition, RAM infected neurons were tested using light stimulations that matched in vivo laser stimulation (15-ms light pulses at 30 Hz). These slices were fixed in 4% PFA in 0.1 M phosphate buffer for three hours at room temperature. Slices containing the mPFC were mounted in PBS and imaged using confocal microscopy (10x, Olympus SpinSR10) to confirm optic fibre placement in the PL. For iDOTS functional isolation, a 100-ms pulse of 405-nm light was delivered to activate terminal PdCO, shunting terminal release from caudal prefrontal fields, immediately followed by a 5-ms pulse of 405-nm light to drive ChR2-expressing inputs from the rostral mPFC. Evoked EPSC amplitudes were normalised within-animal to the maximum recorded amplitude to minimise variance arising from differential viral expression.

### Tissue Processing

Animals were anaesthetised with isoflurane and were transcardially perfused with 4% PFA, then brains were extracted and placed into 4% PFA for three hours at room temperature. Brain tissue was then washed with 0.1 M PBS to remove excess PFA. The brains were then coronally sectioned at a thickness of 100 μm and stained with the Sytox orange nucleic acid stain (Invitrogen) in 0.9% saline (1:10000) or nuclear marker 4′,6-diamidino-2-phenylindole (DAPI) in 0.9% saline (1:5000). Slices at every 200 μm were mounted in PBS and imaged using confocal microscopy (10x, Olympus SpinSR10) at a standardised laser power and exposure time for each channel. Acute slices that were used for electrophysiological recordings were subsequently fixed in 4% paraformaldehyde (PFA) in 0.1 M phosphate buffer for three hours at room temperature. Sections were then washed with 0.1 M phosphate buffered saline (PBS) to remove excess PFA and stained with the nuclear marker 4′,6-diamidino-2-phenylindole (DAPI) in 0.9% saline (1:5000). Slices containing the mPFC and/or BLA were mounted in PBS and imaged using confocal microscopy (10x, Olympus SpinSR10) at a standardised laser power and exposure time for each channel.

For the investigation of overlap between RAM and c-Fos populations in the mPFC, animals were sacrificed two hours post the retrieval session, while off doxycycline feed, to balance allowing enough time for RAM expression while being closest to the peak of c-Fos expression. Brains were perfused and washed according to the protocols described above, then brain sections were stained with primary antibody guinea pig anti-c-Fos (1:10000, Synaptic Systems) at room temperature for 48 h. Then, sections were incubated with Alexa Fluor 647 anti-guinea pig (1:1000, Invitrogen) secondary antibody for 24 h at room temperature. Brain sections were stained with the nuclear marker 4′,6-diamidino-2-phenylindole (DAPI) in 0.9% saline (1:5000) and imaged using confocal microscopy (10x, Olympus SpinSR10).

### Image analysis

Confocal images were produced by flattening z-stacks (3 x 15 μm intervals) to create a maximum projection using ImageJ software. Images of sections were mapped to the Allen coronal reference atlas (Allen Institute for Brain Science, 2008), using the Big Warp ImageJ plugin (BigDataViewer) and in-house adaptations of atlas sections. For cell segmentation and quantification, images were filtered (median, radius = 5), background subtracted (rolling ball, radius = 100) and locally thresholded (Bernsen, radius = 25) using ImageJ. Cell segmentation of thresholded images was performed using CellProfiler (4.0.7) and cells were assigned to a mPFC sub-region and layer by location within the corresponding atlas image. Brain sections with no RAM-driven fluorophore expression were excluded from analysis. For calculating patterns in active cells across the rostro-caudal axis, sections were grouped based on the position in the anterior-posterior plane relative to Bregma. The rostral mPFC was defined as those sections between +2.8 and +2.4 mm Bregma, demarcated by the olfactory bulbs being completely discontinuous with the cortex. The intermediate mPFC was defined as those sections between +2.3 and +2.0 mm Bregma, demarcated by the emergence of the forceps minor fibre bundle. Finally, the caudal mPFC was defined as sections between +1.9 and +1.4 mm Bregma, delimited by the initiation of the corpus callosum amalgamation. These segregations along the rostro-caudal plane were designed to maximise the chance of an even number of sections in each rostro-caudal category. For the quantification of GAD67-EGFP, c-Fos and RAM-positive cells, thresholded images were subtracted in ImageJ (image calculator, AND) to reveal cells that were co-labelled. These positive cells were quantified as a percentage of the total RAM and c-Fos ensemble.

For topographical mapping of efferent and afferent projections along the rostro-caudal axis of the mPFC, serial coronal sections were collected at 200 µm intervals and registered to the Allen Mouse Brain Atlas framework. For anterograde tracing experiments, axonal fluorescence intensity for each unique fluorophore—representing efferent fibres originating from the rostral, intermediate, and caudal mPFC—was quantified within the segmented anatomical boundaries of all regions across the atlas. To eliminate confounding variance introduced by differential viral expression levels and inherent fluorophore brightness, the total fluorescence across the entire brain was summed for each channel, and individual region values were converted into a relative proportion of this whole-brain total. To isolate the signal from efferent projections traveling away from the target site, fluorescence corresponding to the primary injection site (restricted to the ipsilateral side) was subtracted from this dataset. These proportional values were subsequently area-normalised to provide an area-corrected value for unbiased structural comparison across all atlas regions. Finally, to characterise the spatial distribution of fibre inputs along the rostro-caudal axis of the BLA, these area-normalised fluorescence metrics were plotted across longitudinal Bregma coordinates as a percentage of the total area-normalised proportion value calculated for the entire BLA. For the retrograde tracing experiments, an identical analytical and normalisation pipeline was implemented, with the exception that the automated segmentation and quantification of retrogradely labelled somata was used as the readout instead of axonal fluorescence. The final metric was expressed as the area-normalised proportion of total whole-brain projection cells, excluding any local labelling within the primary mPFC injection sites. To map the input topography of the reverse pathway, the spatial distribution of these prefrontal-projecting neurons within the BLA was plotted across the rostro-caudal axis as a percentage of the total area-normalised projection cell population contained within the entire BLA.

### Statistical analysis

Quantification of freezing behaviour was performed using the automated analysis within the FreezeFrame software (Coulbourn Instruments; 0.5-s bouts). Baseline freezing prior to fear conditioning was calculated as the average time spent freezing during the 60 seconds immediately prior to the first tone presentation. The final freezing value during the learning session was calculated as the average time spent freezing during the final two tone presentations. Likewise, the freezing value for the retrieval test was calculated as the average time spent freezing during the two tone presentations. For *in vivo* optogenetic experiments, freezing values were manually scored to remove the effect of optic fibre movement on automated freezing analysis. For phantom retrieval experiments, the baseline was the 60 seconds immediately prior to the first optogenetic stimulation, which was compared to the average freezing levels during the two 60 second periods after both light stimulations (simulating two tone presentations). For inhibition experiments, freezing levels were quantified during the 10 second tone and/or light stimulations, averaged across two tone presentations. All other data analysis and statistical testing was performed using GraphPad Prism (9.5.1), using one-way or two-way ANOVA tests, correcting for multiple comparisons (Tukey’s or Šídák’s test) where appropriate. Statistical outliers were tested for using the Grubb’s method through GraphPad Prism (α=0.01).

## Supplemental Information

Extended Data Figures 1-10

## Acknowledegments

We would like to thank members of the Sah lab for their input and guidance on aspects of this manuscript. We would also like to thank Yingxi Lin’s lab for providing the RAM construct and Ofer Yizhar for providing the PdCo-Chrimson construct, as well as providing guidance throughout the project. We also thank Andreas Lüthi for providing feedback on the manuscript. We thank Phoebe Mayne and Madison Danalis for their assistance with c-Fos immunostaining and analysis. mage acquisition was performed at the Queensland Brain Institute’s Advanced Microscopy Facility using an Olympus SpinSR10 spinning disk confocal microscope, and the authors gratefully acknowledge their support and assistance in this work. This work was supported by grants from the Australian National Health and Medical Research Council (PS) and the Australian Research Council (PS and RM). JPK is employed by the Queensland Centre for Mental Health Research which receives core funding from Queensland Health.

---

**Extended Data Figure 1.**
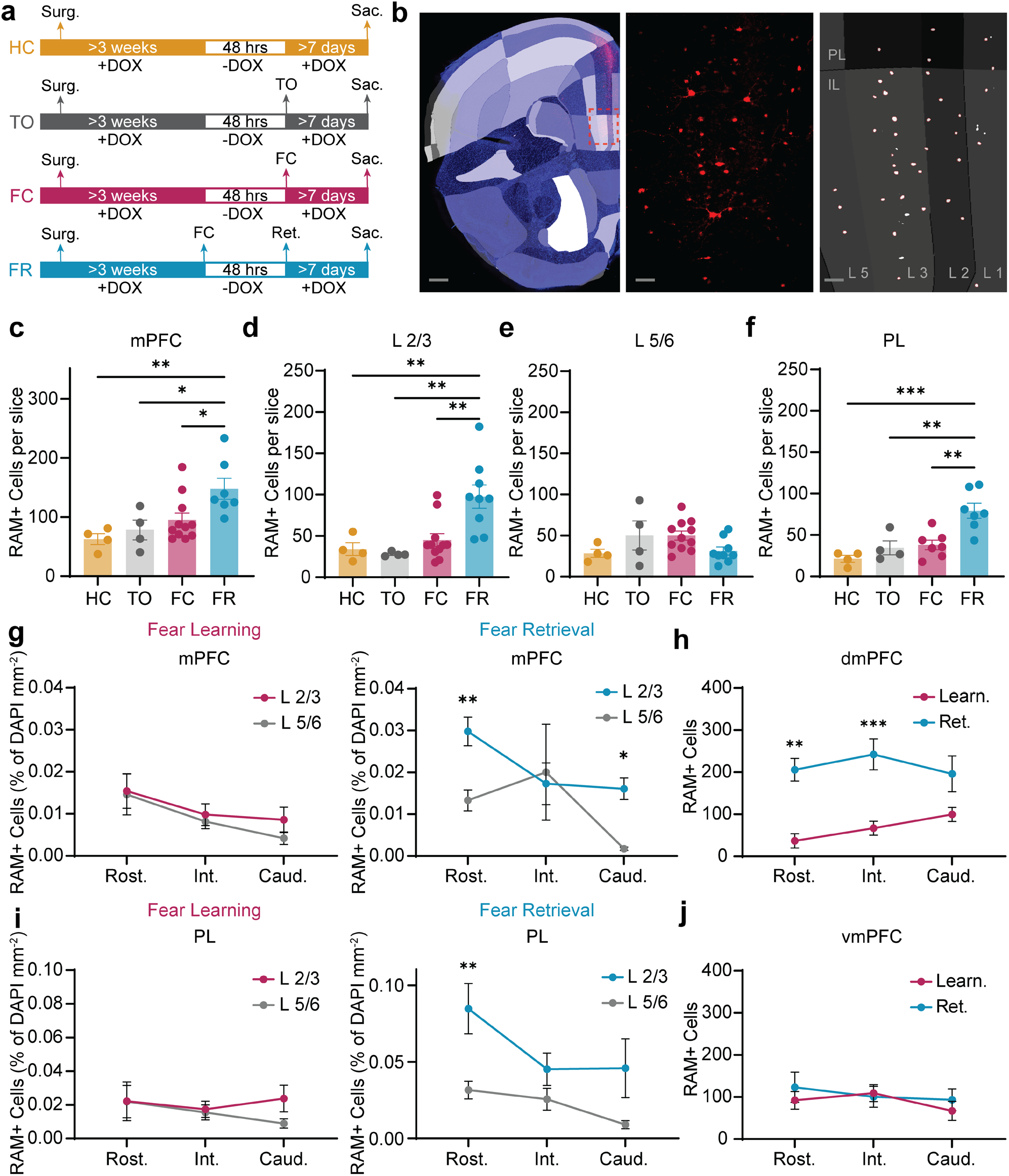
Experimental protocol and changes to the active mPFC ensemble post-learning. **a.** Experimental timelines for the capture of active populations in the mPFC with the RAM system. Animals were kept on doxycycline feed up until 48 h prior to the behaviour of interest, taken off during this window, and returned onto doxycycline feed immediately after the tagged behaviour. At least 7 days was allowed in between behavioural tagging and sacrificing the animal to ensure sufficient time for expression. HC = home cage; TO = tone only; FC = fear learning; FR = fear retrieval. **b.** Image analysis pipeline. Left: Example confocal image of a coronal slice matched to the Allen brain atlas using the BigWarp plugin in ImageJ. Scale bar = 250 μm. Middle: Raw RAM signal of inset (red box) in A. Scale bar = 50 μm. Right: Processed image of RAM signal shown in middle panel, with cell segmentation of RAM labelled cells for analysis. Red outlined cells indicate those cells that were segmented from the thresholded image using CellProfiler. These cells are then quantified based on their position within the overlying atlas image using CellProfiler. Scale bar = 50 μm. **c.** Mean number of active RAM-labelled cells per brain slice across the global mPFC. Fear retrieval induced a significant increase in total active cell count relative to all other groups (P < 0.05, one-way ANOVA with Tukey’s test). **d.** Mean number of active RAM-labelled cells per slice within L2/3 of the mPFC. Fear retrieval drove a significant increase in superficial cell recruitment relative to all other conditions (P < 0.0001, one-way ANOVA with Tukey’s test). **e.** Mean number of active RAM-labelled cells per slice within L5/6 of the mPFC, showing no significant differences across cohorts (P > 0.05, one-way ANOVA). **f.** Mean number of active RAM-labelled cells per brain slice across the PL. Fear retrieval induced a significant increase in total active cell count relative to all other groups (P < 0.01, one-way ANOVA with Tukey’s test). **g.** Total number of active cells within L2/3 and L5/6 (% of DAPI mm^2^) across the rostral, intermediate, and caudal zones of the mPFC. Fear learning (left panel) cells showed no significant differences between layers along the axis (P > 0.05, two-way ANOVA with Šídák’s post-hoc test). Conversely, fear retrieval (right panel) drove a significant increase in active cells within L2/3 of the rostral and caudal mPFC relative to corresponding deep layers (P < 0.01, two-way ANOVA with Šídák’s test). No significant layer-specific differences were found in intermediate or caudal segments (P > 0.05). **h.** Quantification of total RAM labelled cells in the dmPFC along the rostro-caudal axis. Fear retrieval produced a significant increase in active cells in the rostral and intermediate dmPFC, compared to cells labelled during fear learning (rostral: p<0.01, intermediate: p<0.001, two-way ANOVA, Šídák’s test). No significant differences were observed in caudal dmPFC active cells between fear learning and retrieval (p>0.05, two-way ANOVA, Šídák’s test). **i.** Layer-specific count of active cells (% of DAPI mm^2^) restricted within the prelimbic (PL) cortex. Fear learning cells (let panel) showed no layer-specific differences across the rostro-caudal axis (P > 0.05, two-way ANOVA). Fear retrieval (right panel) produced a significant increase in active cells within L2/3 of the rostral PL compared to its deep layers (P < 0.001, two-way ANOVA with Šídák’s test). **j.** Quantification of total RAM labelled cells in the vmPFC along the rostro-caudal axis. No significant differences were found in vmPFC active cells between fear learning and retrieval, across the rostro-caudal axis (p>0.05, two-way ANOVA, Šídák’s test). All data is presented as mean ± SEM.

**Extended Data Figure 2.**
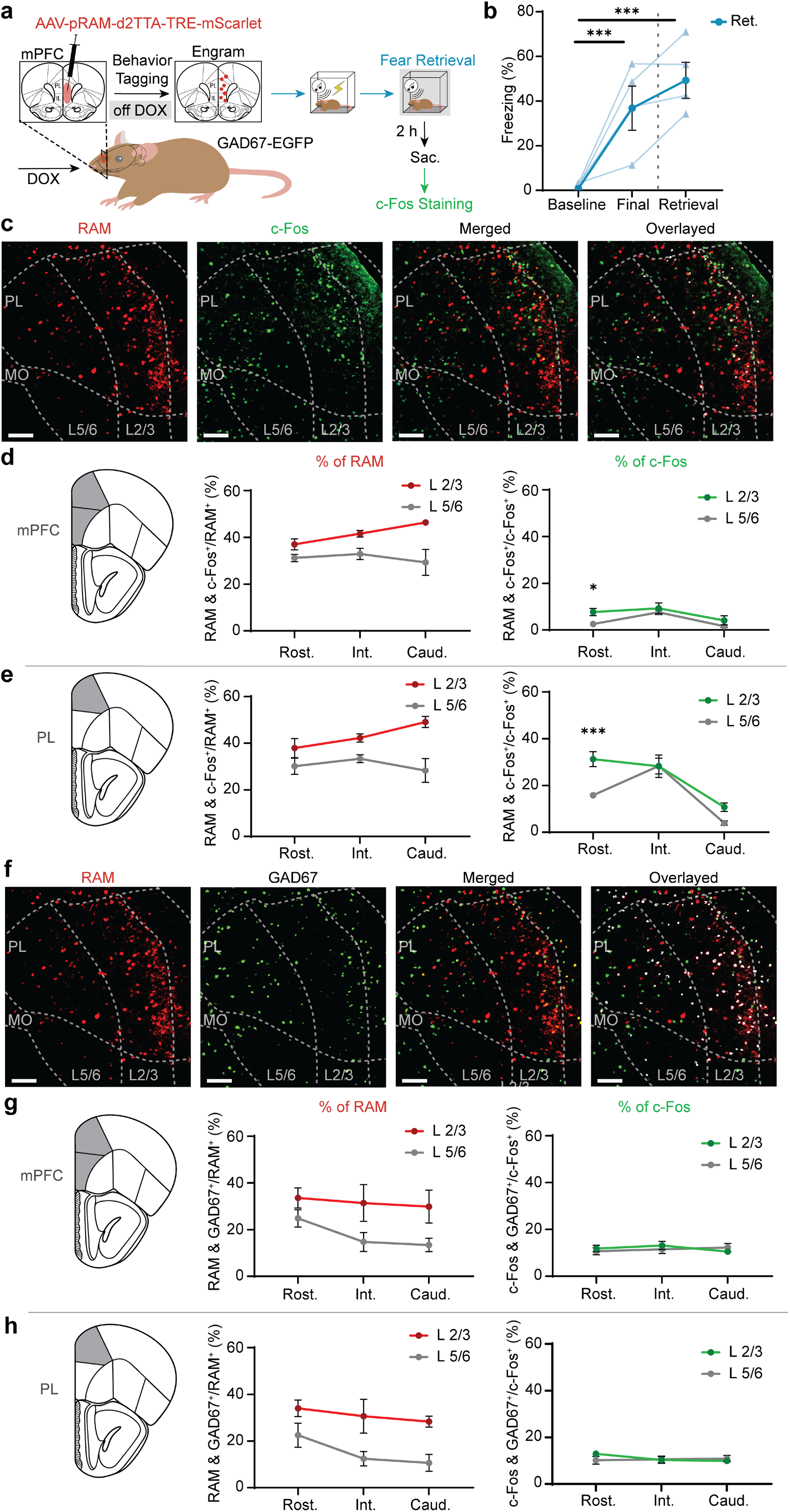
The consolidated mPFC fear engram captures a larger proportion of inhibitory neurons compared to c-Fos immunohistochemistry. **a.** Experimental configuration detailing the viral injection of the activity-dependent RAM system across the mPFC in GAD67-EGFP animals to express mScarlet in neurons active during fear retrieval (n = 4) and the subsequent staining of c-Fos positive neurons using immunohistochemistry. Animals were sacrificed 2 h after fear retrieval to allow labelling of the neurons active within the same behavioural window. **b.** Freezing scores across sessions. Auditory fear conditioning drove significant increases in freezing in the fear retrieval group (baseline vs. final: P < 0.001, two-way ANOVA with Tukey’s multiple comparisons test). Presentation of the CS during the retrieval test drove significant freezing relative to baseline (P < 0.001). **c.** Example confocal images of RAM-labelled (mScarlet+) neurons and c-Fos positive neurons within the mPFC during fear retrieval. The overlayed image (right) shows dual-labelled cells in white that were segmented and used for analysis. Scale bar = 100 μm. **d.** Percentage of RAM/c-Fos+ neurons within L2/3 and L5/6 across the rostral, intermediate, and caudal zones of the mPFC, as a percentage of the total RAM (left) or c-Fos population (right). There were no significant differences in overlap across layers or mPFC rostro-caudal zones (P > 0.05, two-way ANOVA with Šídák’s test). There was a significant higher overlap in L2/3 of the mPFC as a percentage of total c-Fos neurons (P <0.05, two-way ANOVA with Šídák’s test). **e.** Percentage of RAM/c-Fos+ neurons within L2/3 and L5/6 across the rostral, intermediate, and caudal zones of the PL, as a percentage of the total RAM (left) or c-Fos population (right). There were no significant differences in overlap as a percentage of total RAM neurons across layers or mPFC rostro-caudal zones (P > 0.05, two-way ANOVA with Šídák’s test). There was a significant higher overlap in L2/3 of the PL as a percentage of total c-Fos neurons (P <0.001, two-way ANOVA with Šídák’s test). There was also significantly more overlap (% of c-Fos) in the rostral and intermediate PL, compared to the caudal PL (row effect, P <0.0001, two-way ANOVA). **f.** Example confocal images of RAM-labelled (mScarlet+) neurons and GAD67-EGFP positive neurons within the mPFC during fear retrieval. The overlayed image (right) shows dual-labelled cells in white that were segmented and used for analysis. Scale bar = 100 μm. **g.** Percentage of RAM/GAD67+ (left) and c-Fos/GAD67+ (right) neurons within L2/3 and L5/6 across the rostral, intermediate, and caudal zones of the mPFC, as a percentage of the total RAM (left) or c-Fos population (right). There were no significant differences in overlap across layers (P > 0.05, two-way ANOVA with Šídák’s test). There was a significantly larger number of dual-labelled RAM/GAD67+ neurons in L2/3 across all zones of the mPFC (column effect, P <0.01, two-way ANOVA). **h.** Percentage of RAM/GAD67+ (left) and c-Fos/GAD67+ (right) neurons within L2/3 and L5/6 across the rostral, intermediate, and caudal zones of the PL, as a percentage of the total RAM (left) or c-Fos population (right). There were no significant differences in overlap across layers (P > 0.05, two-way ANOVA with Šídák’s test). There was a significantly larger number of dual-labelled RAM/GAD67+ neurons in L2/3 across all zones of the PL (column effect, P <0.001, two-way ANOVA). All data is presented as mean ± SEM.

**Extended Data Figure 3.**
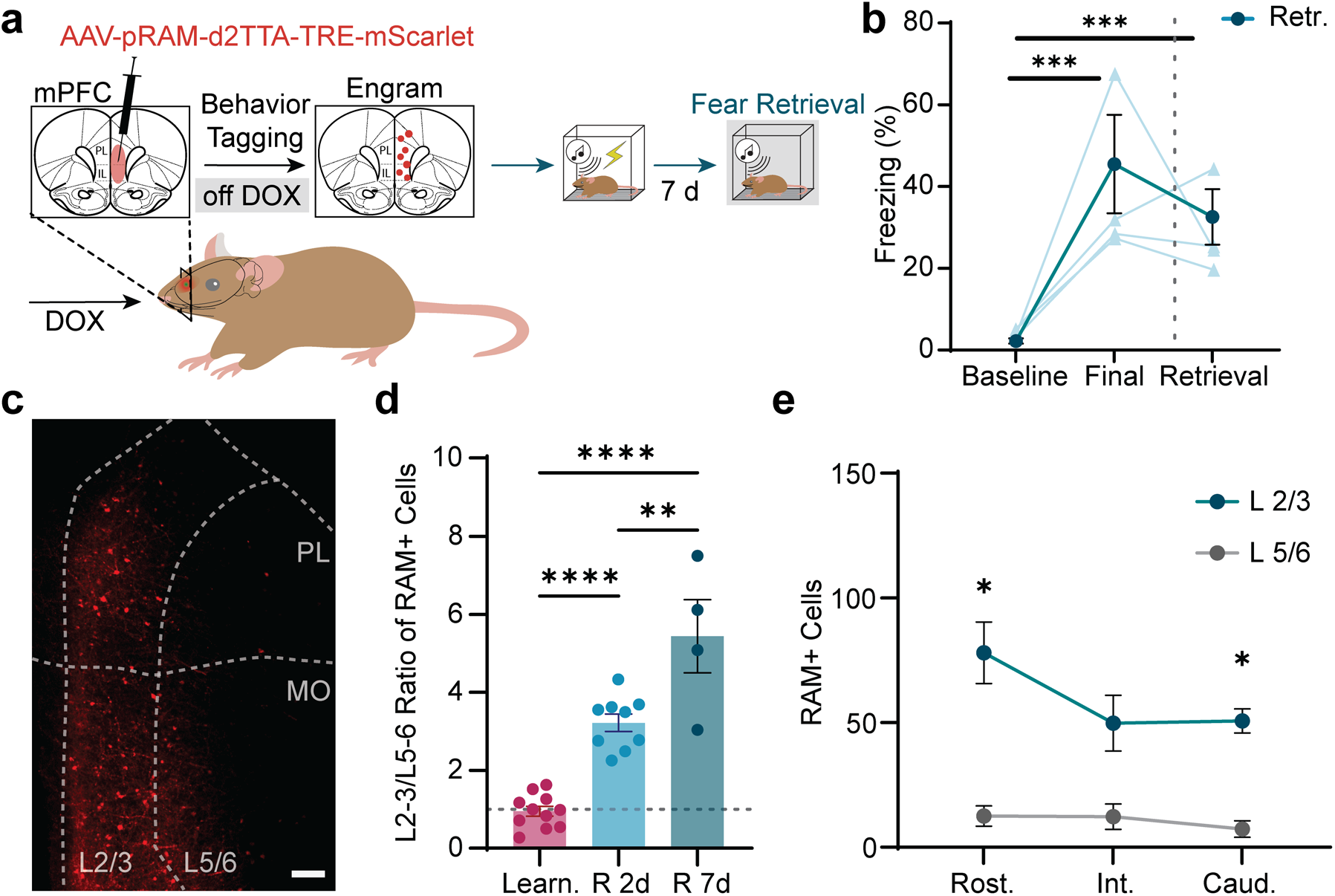
Dominant layer 2/3 activity in the mPFC during the remote retrieval of fear memory. **a.** Experimental configuration detailing the viral injection of the activity-dependent RAM system across the mPFC to express mScarlet in neurons active during a remote fear retrieval test 7 days post fear learning (n = 4). **b.** Freezing scores across sessions. Auditory fear conditioning drove significant increases in freezing in the remote fear retrieval group (baseline vs. final: P < 0.001, two-way ANOVA with Tukey’s multiple comparisons test). Presentation of the CS during the remote retrieval test drove significant freezing relative to baseline (P < 0.001). **c.** Example confocal image of RAM labelled neurons in the mPFC across L2/3 and L5/6 during the remote retrieval of fear. Scale bar = 100 μm. **d.** Superficial-to-deep ratio (L2/3 vs. L5/6) of active cells across the mPFC. Remote fear retrieval (R 7d) drove a significant increase in this ratio compared to fear learning and fear retrieval (R 2d) groups (P < 0.01, one-way ANOVA with Tukey’s test). Data from fear learning and fear retrieval groups reproduced from Fig. 1 for comparison. **e.** Total number of active cells within L2/3 and L5/6 across the rostral, intermediate, and caudal zones of the global mPFC during remote fear retrieval. There was a significant increase in the number of RAM-labelled neurons in L2/3 compared to L5/6 across all mPFC zones (P < 0.05, two-way ANOVA with Šídák’s test). All data is presented as mean ± SEM.

**Extended Data Figure 4.**
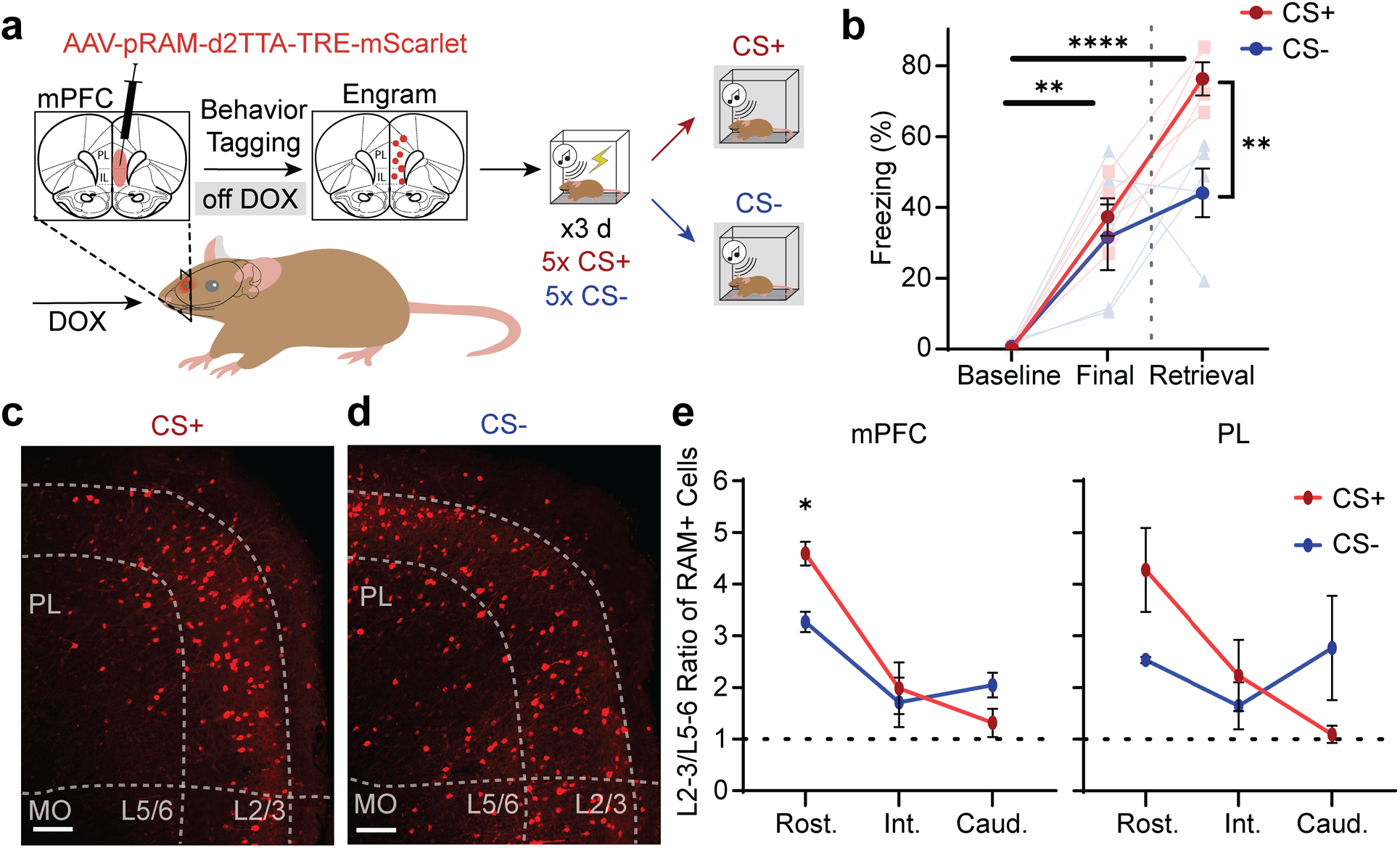
Alterations to the fear engram pattern of activity in the mPFC during differential conditioning. **a.** Experimental configuration detailing the viral injection of the activity-dependent RAM system across the mPFC to express mScarlet in neurons active during the retrieval of differential conditioning learning. Animals underwent 3 consecutive days of fear conditioning with 5 presentations of CS+ (4 kHz) and a CS-(10 kHz), with the CS+ coterminating with a foot-shock. After learning, animals were divided into CS+ (red) and CS-(blue) retrieval groups, whereby the RAM system was used to label the mPFC ensemble active during retrieval of the memory associated with either the CS+ or CS-. **b.** Freezing scores across sessions. Auditory fear conditioning drove significant increases in freezing in both the CS+ and CS-group (baseline vs. final: P < 0.01, two-way ANOVA with Tukey’s test). Presentation of the CS+ during the retrieval test drove significantly greater freezing levels relative to the CS-(P < 0.01). **c.** Example confocal image of RAM labelled neurons in the mPFC across L2/3 and L5/6 during the retrieval of fear memory associated with the CS+. Scale bar = 100 μm. **d.** Example confocal image of RAM labelled neurons in the mPFC across L2/3 and L5/6 during the retrieval of fear memory associated with the CS-. Scale bar = 100 μm. **e.** Superficial-to-deep ratio (L2/3 vs. L5/6) of active cells across the mPFC. The retrieval of the memory associated with the CS+ drove a higher superficial-to-deep layer ratio of active cells in the rostral mPFC compared to CS-retrieval (P < 0.05, two-way ANOVA with Tukey’s test). There was a significant difference in the distribution of active cells in superficial to deep layers across the rostro-caudal zones of the mPFC (interaction effect, P < 0.05, two-way ANOVA). **f.** Superficial-to-deep ratio (L2/3 vs. L5/6) of active cells across the PL. There was a significant difference in the distribution of active cells in superficial to deep layers across the rostro-caudal zones of the mPFC (interaction effect, P < 0.05, two-way ANOVA).

**Extended Data Figure 5.**
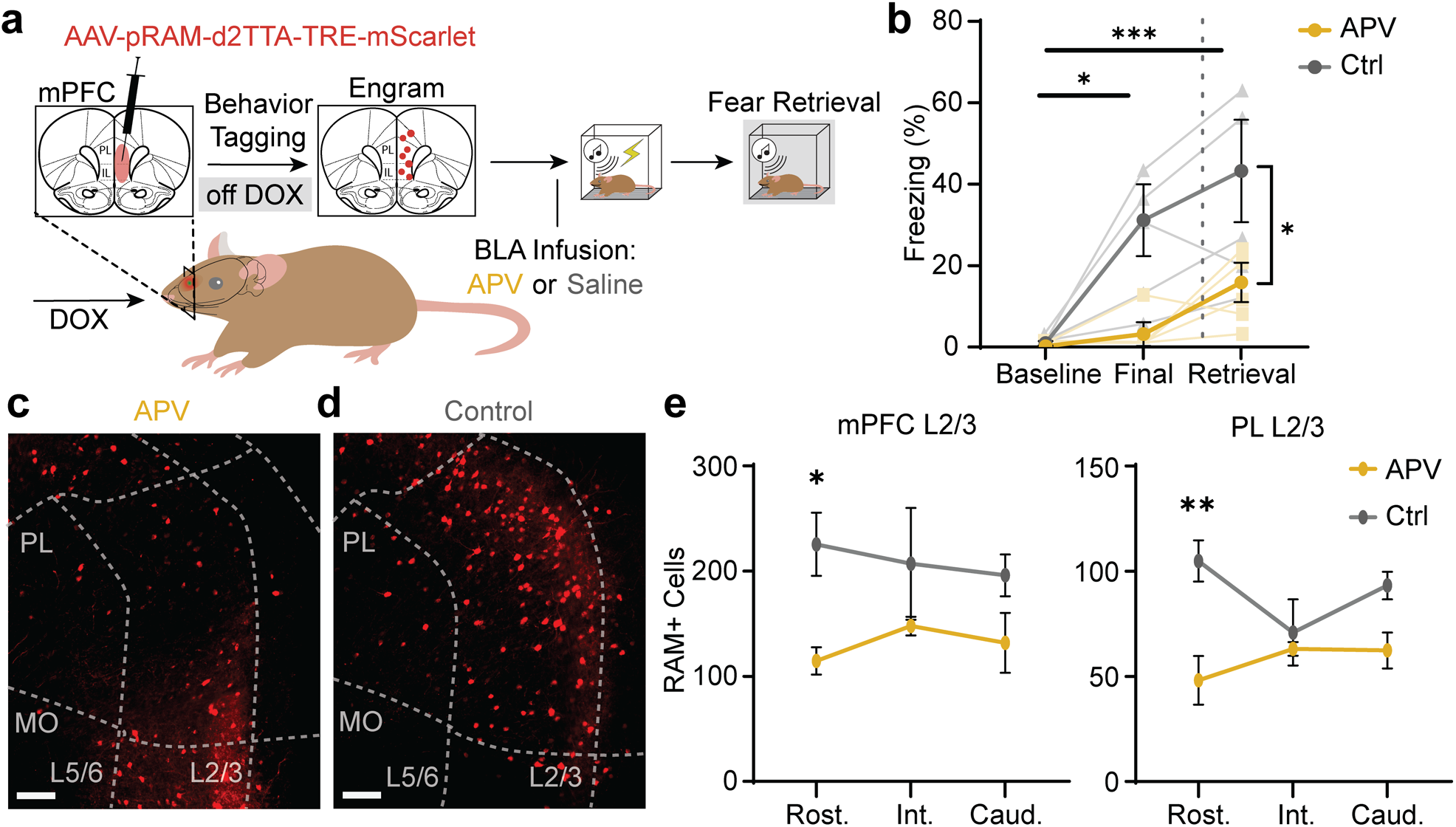
Disruption of fear learning with APV alters the fear engram pattern in the mPFC during subsequent retrieval. **a.** Experimental configuration detailing the viral injection of the activity-dependent RAM system across the mPFC to express mScarlet in neurons active during a fear retrieval test after fear learning has been disrupted using a BLA infusion of APV. Animals received a BLA infusion of APV (yellow) or saline (control, grey) 30 minutes prior to fear conditioning. The RAM system was used to capture active cells in the mPFC during a subsequent retrieval. **b.** Freezing scores across sessions. Auditory fear conditioning drove significant increases in freezing in the control group (baseline vs. final: P < 0.05, two-way ANOVA with Tukey’s test), but not in the APV group (P > 0.05). Presentation of the CS during the retrieval test drove significantly greater freezing levels in the control group relative to the APV group (P < 0.05, two-way ANOVA with Tukey’s test). **c.** Example confocal image of RAM labelled neurons in the mPFC across L2/3 and L5/6 during the retrieval of fear memory in the APV group. Scale bar = 100 μm. **d.** Example confocal image of RAM labelled neurons in the mPFC across L2/3 and L5/6 during the retrieval of fear memory in the control (saline) group. Scale bar = 100 μm. **e.** Total number of active cells within L2/3 across the rostral, intermediate, and caudal zones of the mPFC in APV and control groups. There was a significant decrease in the number of RAM-labelled neurons in L2/3 of the rostral mPFC in the APV group, compared to the control group (P < 0.05, two-way ANOVA with Šídák’s test). **f.** Total number of active cells within L2/3 across the rostral, intermediate, and caudal zones of the PL in APV and control groups. There was a significant decrease in the number of RAM-labelled neurons in L2/3 of the rostral mPFC in the APV group, compared to the control group (P < 0.01, two-way ANOVA with Šídák’s test). All data is presented as mean ± SEM.

**Extended Data Figure 6.**
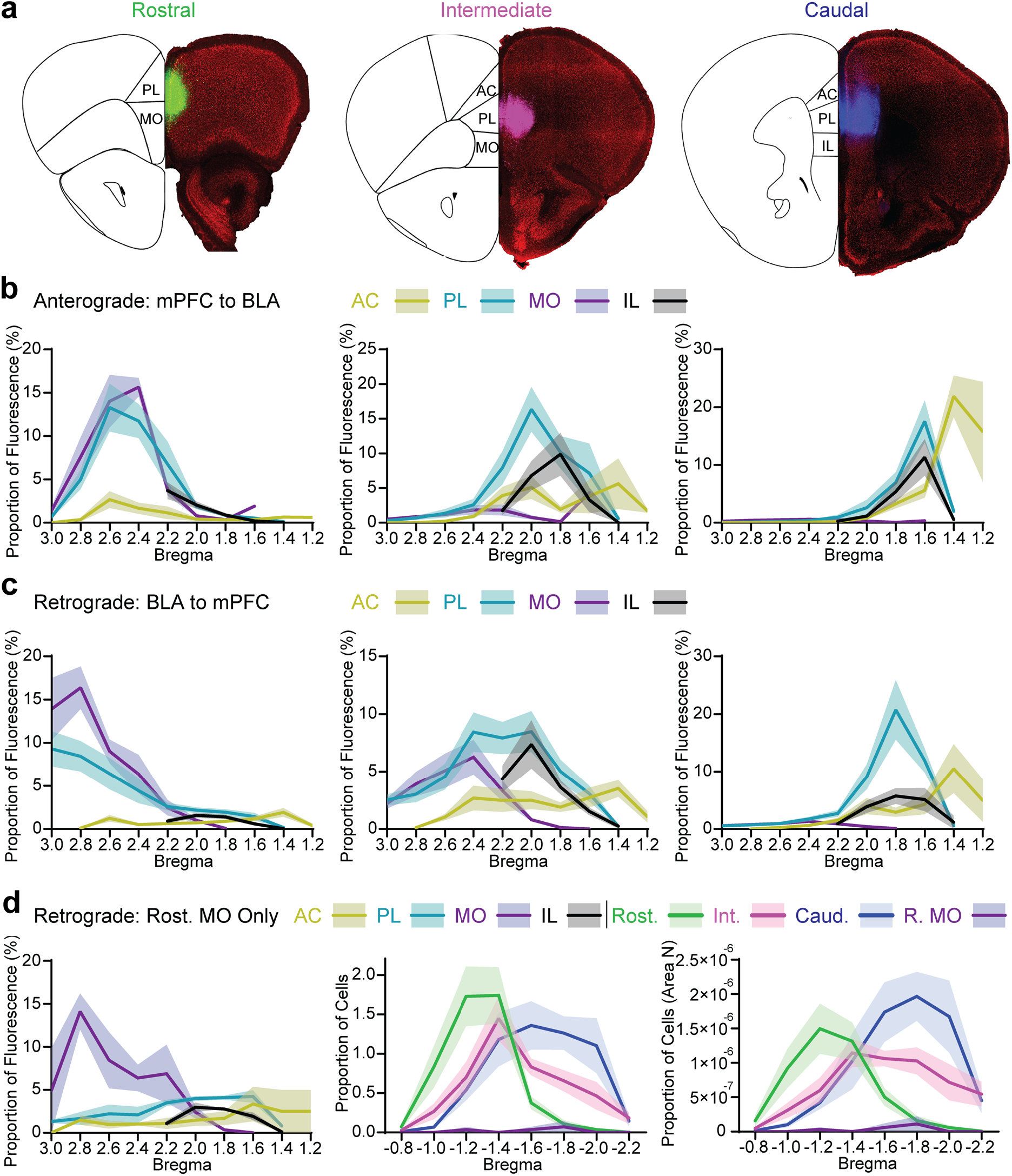
Injection site profiles of tracing of reciprocal connections between the mPFC and BLA. **a.** Example confocal images of viral expression of three retrograde or anterograde tracers with unique fluorophores (e.g. GFP, pemiRFP-670-N1, mTagBFP2) in the rostral (left), intermediate (middle) or caudal (right) mPFC zones. It was ensured that each injection site contained considerable expression in the PL across all mPFC zones. **b.** Mean injection site expression profiles for anterograde tracing of projections from the mPFC to the BLA across the rostral (left), intermediate (middle) or caudal (right) mPFC zones (n = 7 mice). Contributions from the different sub-regions of the mPFC are represented by various colours. **c.** Mean injection site expression profiles for retrograde tracing of projections from the mPFC to the BLA across the rostral (left), intermediate (middle) or caudal (right) mPFC zones (n = 12 mice). Contributions from the different sub-regions of the mPFC are represented by various colours. **d.** Left: Mean injection site profiles for a sub-set of retrograde tracing of projections from the BLA to the rostral mPFC in which expression was isolated to the rostral MO (n = 2 mice, P < 0.05, two-way ANOVA with Tukey’s test). Middle: Quantification of the proportion of total cells in the BLA projecting to rostral MO, as well as the rostral, intermediate and caudal zones of the mPFC. An injection site limited to the rostral MO produced significantly fewer projection cells in the BLA, compared to those projecting to the rostral, intermediate and caudal zones of the mPFC (P < 0.01, two-way ANOVA with Tukey’s test). Right: Quantification of the area normalised (Area N) proportion of total cells in the BLA projecting to rostral MO, as well as the rostral, intermediate and caudal zones of the mPFC. An injection site limited to the rostral MO produced significantly fewer projection cells in the BLA, compared to those projecting to the rostral, intermediate and caudal zones of the mPFC (P < 0.01, two-way ANOVA with Tukey’s test). All data is presented as mean ± SEM.

**Extended Data Figure 7.**
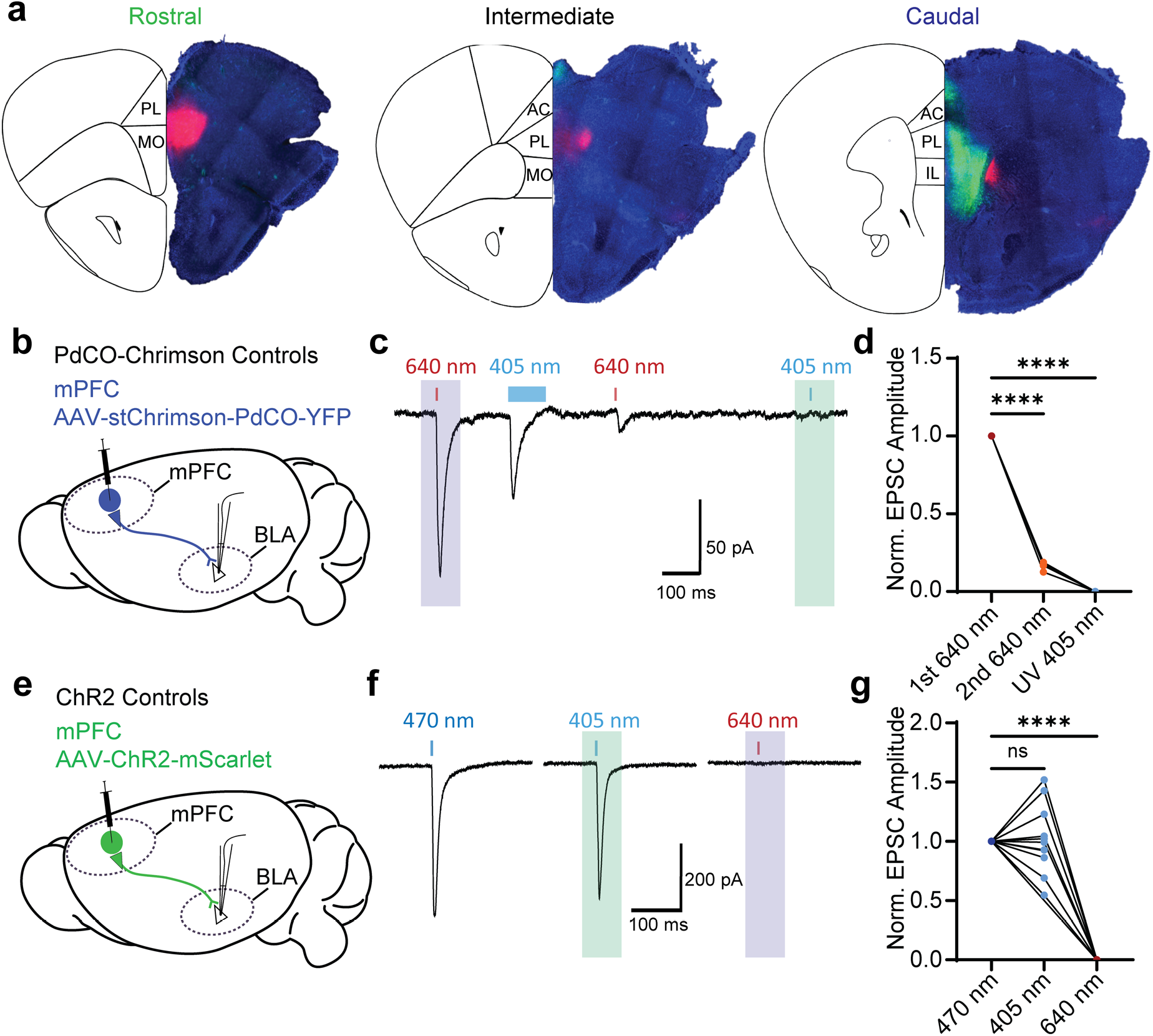
independent dual-opsin terminal stimulation (iDOTS) controls. Viral constructs used in the iDOTS system - Chr2-mScarlet and stChrimson-PdCO-YFP were injected into the rostral and caudal mPFC respectively. It was ensured that each injection site contained considerable expression in the PL and that the two injection sites were distinct with the intermediate mPFC largely devoid of expression of either construct. **a.** Example confocal images of expression of Chr2-mScarlet and stChrimson-PdCO-YFP in the rostral (left), intermediate (middle) or caudal (right) mPFC zones. **b.** Experimental configuration for the Chrimson-PdCO control experiments, detailing the viral injection of stChrimson-PdCO into the mPFC and subsequent electrophysiological recordings from the BLA in acute brain slices. The aim of the control experiments was to ascertain whether combining Chrimson and PdCO could limit the activation of presynaptic terminals in the BLA to red wavelengths (640 nm), while eliminating the smaller activation of Chrimson at blue wavelengths. **c.** Representative EPSC trace recorded from a BLA neuron in a Chrimson-PdCO control animal. Stimulating Chrimson-PdCO with a 640 nm light pulse (5 ms) evokes a large monosynaptic response in the BLA, which is greatly reduced during a subsequent 640 nm light pulse (5 ms, 2nd 640 nm), following PdCO activation with 405 nm light (100 ms). Subsequent stimulation of the PdCO-inhibited terminals with a 405 nm pulse (5 ms, desired wavelength for ChR2 activation) evokes no postsynaptic response. Shaded boxes designate the analysis windows for isolated inputs from the caudal (blue) and rostral (green) mPFC in combined iDOTS experiments. All recordings were performed at –60 mV. **d.** Quantification of normalised EPSC responses during the initial 640 nm pulse (1st 640 nm), second 640 nm pulse post-PdCO activation (2nd 640 nm) and final UV 405 nm pulse (n = 4 cells). Activation of PdCO produced a significant reduction in the response to the 2nd 640 nm pulse (P < 0.0001, one-way ANOVA with Tukey’s test) and a further reduction to zero postsynaptic response when stimulating with a UV 405 nm pulse (P < 0.0001, one-way ANOVA with Tukey’s test). **e.** Experimental configuration for the ChR2 control experiments, detailing the viral injection of ChR2 into the mPFC and subsequent electrophysiological recordings from the BLA in acute brain slices. The aim of the control experiments was to ascertain whether stimulating ChR2 with 405 nm light would produce equitable postsynaptic responses (allowing the use of a wavelength further from the peak of Chrimson) and that stimulating ChR2 terminals with 640 nm light does not evoke postsynaptic responses, even at high expression levels. **f.** Representative EPSC traces recorded from a single BLA neuron in a ChR2 control animal. Stimulating ChR2-expressing mPFC fibres in the BLA with short 470 nm or 405 nm light pulses (5 ms) evoked similar postsynaptic responses in BLA neurons. Stimulating the same terminals with a short 640 nm pulse (5 ms) evoked no postsynaptic response. **g.** Quantification of normalised EPSC responses during the 470 nm pulse, 405 nm and 640 nm pulse (n = 16 cells). There were no significant differences in normalised EPSC responses to 470 nm and 405 nm light pulses (P > 0.05, one-way ANOVA with Tukey’s test). There was a significant reduction to zero postsynaptic response when stimulating with a 640 nm pulse (P < 0.0001, one-way ANOVA with Tukey’s test).

**Extended Data Figure 8.**
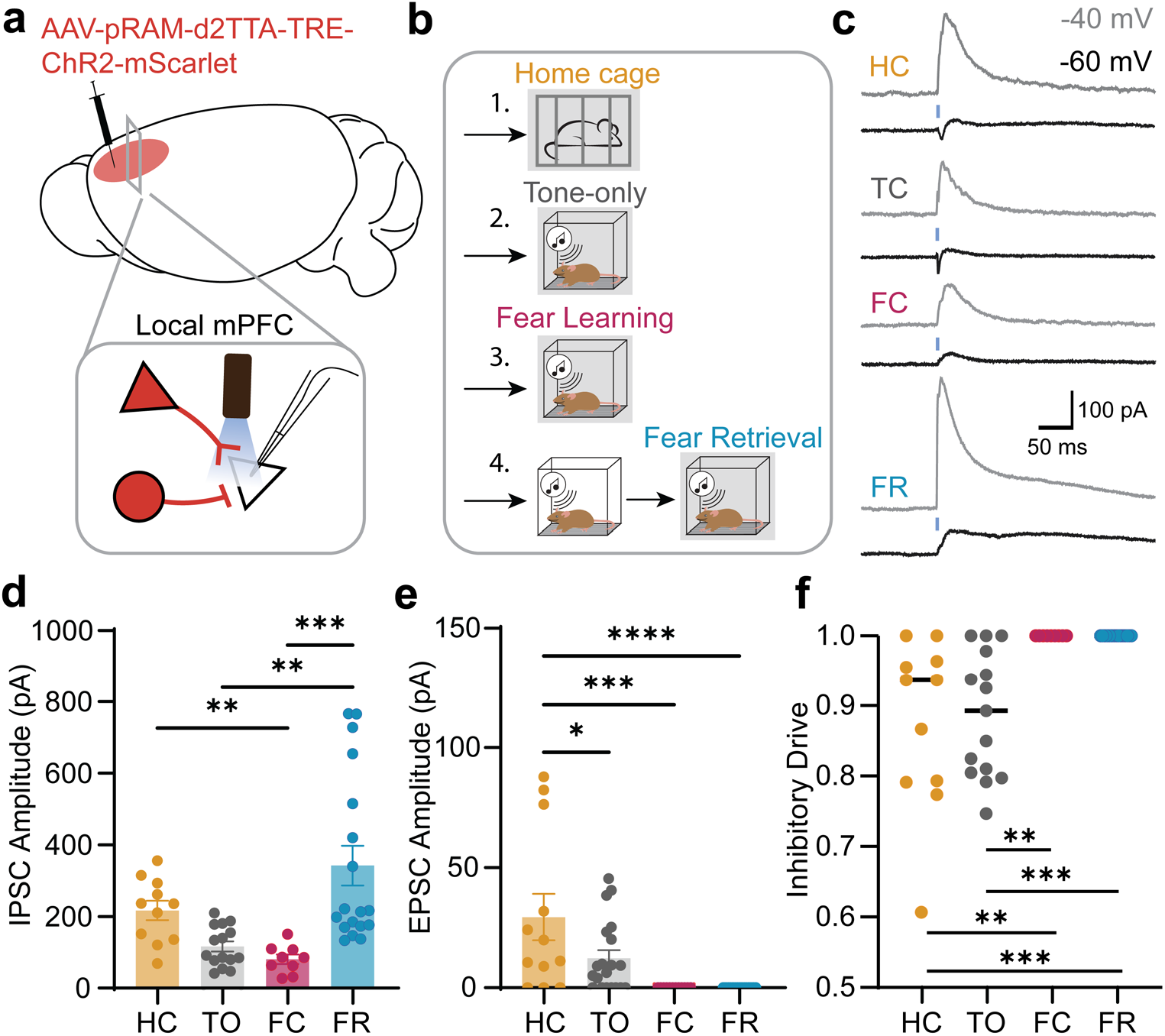
Consolidated fear memory features dominant inhibition in L2/3 of the mPFC. **a.** Experimental strategy for activity-dependent local circuitry mapping. The RAM system was injected into the mPFC to drive ChR2 and mScarlet expression in cells active during specific behavioural windows. At least one-week post-tagging, acute brain slices were taken and whole-cell recordings were made from non-labelled (non-engram) neurons in L2/3 of the mPFC during optical reactivation of prefrontal axon terminals. **b.** The RAM system was used to capture active neuronal ensembles in the mPFC under four different behavioural conditions: home cage (HC, orange, n = 4), tone-only (TO, grey, n = 4), fear learning (FC, purple, n = 5) and fear retrieval (FR, blue, n = 5). **c.** Representative light-evoked responses from non-engram mPFC neurons when mPFC active ensemble axon terminals were optically stimulated across all behavioural conditions. Blue bars denote the 5 ms, 470 nm light pulse. Inhibitory responses were recorded at-40 mV (grey), while excitatory responses were recorded at –60 mV (black). **d.** Quantification of IPSC response amplitudes across all conditions, recorded at-40 mV. Fear retrieval produced significantly larger IPSC amplitudes in non-labelled layer 2/3 neurons, compared to fear learning (p<0.001) and tone-only conditions (p<0.01, one-way ANOVA, Tukey’s test). **e.** Quantification of EPSC response amplitudes across all conditions, recorded at-60 mV. Home cage conditions produced significantly larger EPSC amplitudes in non-labelled layer 2/3 neurons, compared to fear learning (p<0.001), fear retrieval (p<0.001) and tone-only conditions (p<0.05, one-way ANOVA, Tukey’s test). **f.** Quantification of the balance of inhibitory drive (ratio of IPSC and EPSC amplitude) for non-labelled L2/3 neurons across all conditions. Home cage and tone-only conditions produced significantly lower inhibitory drive, compared to fear learning (p<0.01) and fear retrieval conditions (p<0.01, one-way ANOVA, Tukey’s test). All data are presented as mean ± SEM.

**Extended Data Figure 9.**
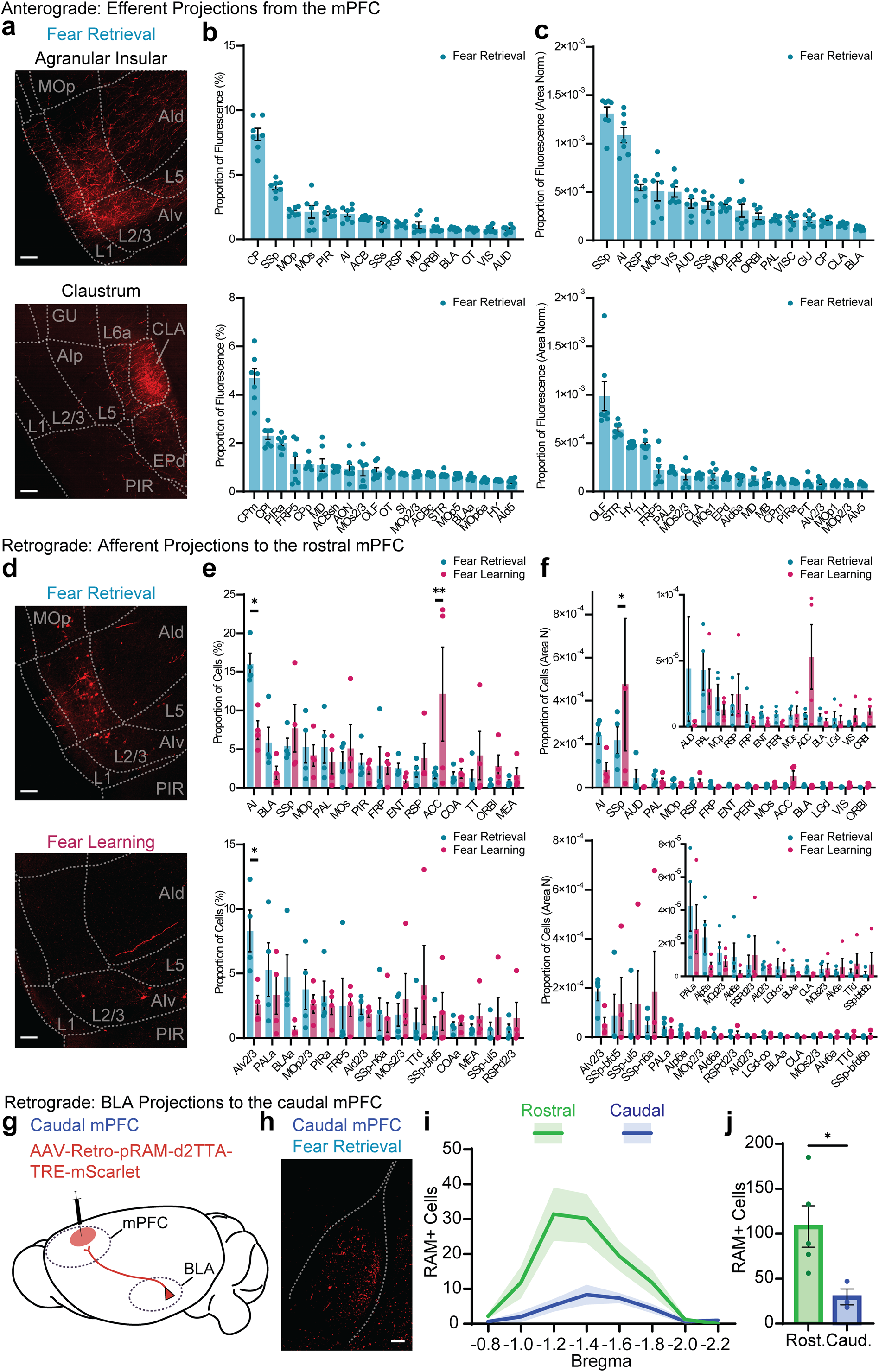
Full brain mapping of the mPFC fear engram. **a.** Example confocal images of axon terminals in the agranular insular (AI, top) and claustrum (CLA, below), originating from mPFC fear retrieval engram neurons. MOp = primary motor area, AId = dorsal agranular insular, AIv = ventral agranular insular, AIp = posterior agranular insular, GU = gustatory areas, EPd = dorsal endopiriform nucleus, PIR = piriform area. Scale bar = 100 μm. **b.** Quantification of the proportion of total brain fluorescence (%) of fear retrieval engram axon terminals across the highest contributing major (top) and minor (below) brain atlas regions (n = 7 mice). **c.** Quantification of the area normalised proportion of total brain fluorescence of fear retrieval engram axon terminals across the highest contributing major (top) and minor (below) brain atlas regions (n = 7 mice). **d.** Example confocal images of retrogradely labelled neurons in the agranular insular projecting to the mPFC, active during fear retrieval (blue, top) or fear learning (purple, below). MOp = primary motor area, AId = dorsal agranular insular, AIv = ventral agranular insular, PIR = piriform area. Scale bar = 100 μm. **e.** Quantification of the proportion of total retrogradely labelled cells (%) of the active neuronal population during fear retrieval (blue, n = 4 mice) and fear learning (purple, n = 4 mice) across the highest contributing major (top) and minor (below) brain atlas regions. Top contributing regions were sorted according to the fear retrieval engram population. **f.** Quantification of the area normalised proportion of total retrogradely labelled cells of the active neuronal population during fear retrieval (blue, n = 4 mice) and fear learning (purple, n = 4 mice) across the highest contributing major (top) and minor (below) brain atlas regions. Inset: rescaled graph excluding the top contributing regions for visualisation purposes. Top contributing regions were sorted according to the fear retrieval engram population. **g.** Experimental strategy for activity-dependent retrograde input mapping. Retro-RAM was virally delivered into the caudal mPFC to express mScarlet in active afferent projection neurons during fear retrieval (n = 3). **h.** Representative confocal image of retrogradely labelled active (RAM+) cells within the BLA targeting the caudal mPFC during fear retrieval. Scale bar = 100 µm. **i.** Distribution profile of RAM-labelled cells projecting to the rostral (green, n = 5, raw data from Fig. 6) and caudal mPFC (blue, n = 3) across the rostro-caudal axis of the BLA. **j.** Quantification of total number of retrogradely labelled RAM+ cells projecting to the rostral and caudal mPFC during fear retrieval. There were significantly more active cells projecting to the rostral mPFC than the caudal mPFC (P < 0.05, unpaired t test). All data are presented as mean ± SEM.

**Extended Data Figure 10.**
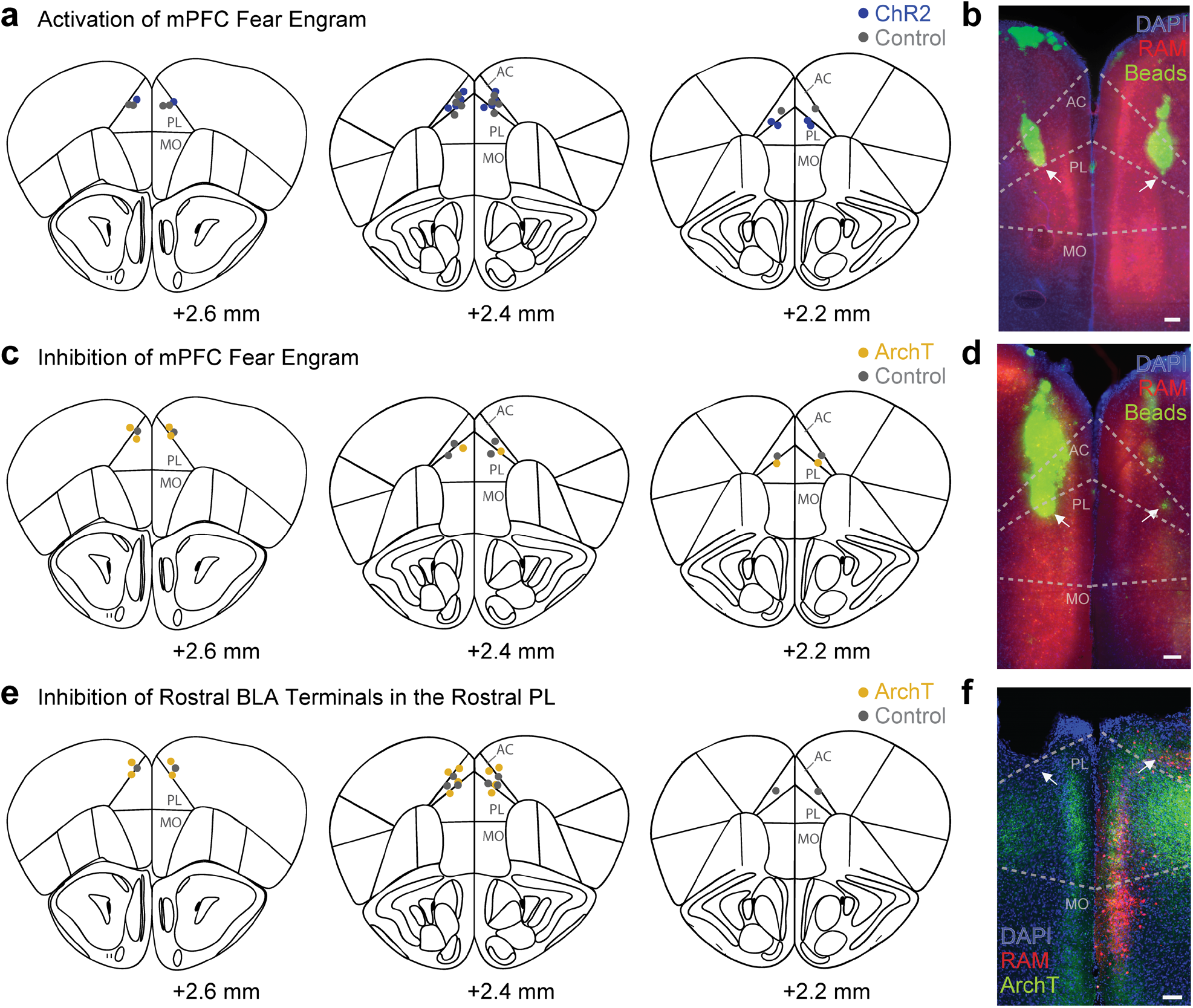
Optic fibre placements and viral expression for optogenetic manipulation experiments. **a.** Schematic representations of coronal mouse brain slices (+2.6 mm, +2.4 mm, and +2.2 mm relative to Bregma) displaying verified bilateral optical fibre tip locations within the prelimbic cortex (PL) for ChR2 (blue dots) and control (mScarlet, grey) groups during optogenetic reactivation of fear retrieval engrams. **b.** Representative image of the mPFC showing engram expression (RAM, red), fluorescent beads demarcating fibre placement tracks (green, white arrows), and nuclear stain (DAPI, blue). **c.** Schematic representations of coronal mouse brain slices (+2.6 mm, +2.4 mm, and +2.2 mm relative to Bregma) displaying verified bilateral optical fibre tip locations within the prelimbic cortex (PL) for ArchT (yellow) and control (mScarlet, grey) groups during optogenetic inhibition of fear retrieval engrams. **d.** Representative image of the mPFC showing engram expression (RAM, red), fluorescent beads demarcating fibre placement tracks (green, white arrows), and nuclear stain (DAPI, blue). **e.** Schematic representations of coronal mouse brain slices (+2.6 mm, +2.4 mm, and +2.2 mm relative to Bregma) displaying verified bilateral optical fibre tip locations within the prelimbic cortex (PL) for ArchT (yellow) and control (mScarlet, grey) groups during optogenetic inhibition of rostral BLA terminals in the rostral PL. **f.** Representative confocal image showing terminal innervation of ArchT (green), RAM expression (red) and DAPI (blue) in the mPFC. White arrows denote fibre insertion paths.

## References

1 Dudai, Y. The neurobiology of consolidations, or, how stable is the engram? Annu Rev Psychol 55, 51–86, doi:10.1146/annurev.psych.55.090902.142050 (2004).

2 Takehara-Nishiuchi, K. Neuronal ensemble dynamics in associative learning. Curr Opin Neurobiol 73, 102530, doi:10.1016/j.conb.2022.102530 (2022).

3 de Sousa, A. F., Chowdhury, A. & Silva, A. J. Dimensions and mechanisms of memory organization. Neuron 109, 2649–2662, doi:10.1016/j.neuron.2021.06.014 (2021).

4 Goto, A. et al. Stepwise synaptic plasticity events drive the early phase of memory consolidation. Science 374, 857–863, doi:10.1126/science.abj9195 (2021).

5 Roy, D. S. et al. Brain-wide mapping reveals that engrams for a single memory are distributed across multiple brain regions. Nature Communications 13, 1799, doi:10.1038/s41467-022-29384-4 (2022).

6 Tomé, D. F. et al. Dynamic and selective engrams emerge with memory consolidation. Nature Neuroscience 27, 561–572, doi:10.1038/s41593-023-01551-w (2024).

7 Semon, R. W. The mneme. (G. Allen & Unwin Limited, 1921).

8 Semon, R. W. Die Mneme als erhaltendes Prinzip im Wechsel des organischen Geschehens. (Engelmann, 1920).

9 Semon, R. W. Mnemic psychology (G. Allen & Unwin Limited, 1923).

10 Tonegawa, S., Liu, X., Ramirez, S. & Redondo, R. Memory Engram Cells Have Come of Age. Neuron 87, 918–931, doi:10.1016/j.neuron.2015.08.002 (2015).

11 Kitamura, T. et al. Engrams and circuits crucial for systems consolidation of a memory. Science 356, 73–78, doi:10.1126/science.aam6808 (2017).

12 Liu, X. et al. Optogenetic stimulation of a hippocampal engram activates fear memory recall. Nature 484, 381–385, doi:10.1038/nature11028 (2012).

13 Tayler, Kaycie K., Tanaka, Kazumasa Z., Reijmers, Leon G. & Wiltgen, Brian J. Reactivation of Neural Ensembles during the Retrieval of Recent and Remote Memory. Current Biology 23, 99–106, doi:10.1016/j.cub.2012.11.019 (2013).

14 Reijmers, L. G., Perkins, B. L., Matsuo, N. & Mayford, M. Localization of a Stable Neural Correlate of Associative Memory. Science 317, 1230–1233, doi:10.1126/science.1143839 (2007).

15 Denny, Christine A. et al. Hippocampal Memory Traces Are Differentially Modulated by Experience, Time, and Adult Neurogenesis. Neuron 83, 189–201, doi:10.1016/j.neuron.2014.05.018 (2014).

16 Cho, H.-Y. et al. Turnover of fear engram cells by repeated experience. Current Biology 31, 5450–5461.e5454, doi:10.1016/j.cub.2021.10.004 (2021).

17 DeNardo, L. A. et al. Temporal evolution of cortical ensembles promoting remote memory retrieval. Nat Neurosci 22, 460–469, doi:10.1038/s41593-018-0318-7 (2019).

18 Yizhar, O. & Klavir, O. Reciprocal amygdala-prefrontal interactions in learning. Curr Opin Neurobiol 52, 149–155, doi:10.1016/j.conb.2018.06.006 (2018).

19 Marek, R., Sun, Y. & Sah, P. Neural circuits for a top-down control of fear and extinction. Psychopharmacology (Berl) 236, 313–320, doi:10.1007/s00213-018-5033-2 (2019).

20 Sierra-Mercado, D., Jr., Corcoran, K. A., Lebron-Milad, K. & Quirk, G. J. Inactivation of the ventromedial prefrontal cortex reduces expression of conditioned fear and impairs subsequent recall of extinction. Eur J Neurosci 24, 1751–1758, doi:10.1111/j.1460-9568.2006.05014.x (2006).

21 Corcoran, K. A. & Quirk, G. J. Activity in prelimbic cortex is necessary for the expression of learned, but not innate, fears. J Neurosci 27, 840–844, doi:10.1523/JNEUROSCI.5327-06.2007 (2007).

22 Do-Monte, F. H., Quinones-Laracuente, K. & Quirk, G. J. A temporal shift in the circuits mediating retrieval of fear memory. Nature 519, 460–463, doi:10.1038/nature14030 (2015).

23 Vidal-Gonzalez, I., Vidal-Gonzalez, B., Rauch, S. L. & Quirk, G. J. Microstimulation reveals opposing influences of prelimbic and infralimbic cortex on the expression of conditioned fear. Learn Mem 13, 728–733, doi:10.1101/lm.306106 (2006).

24 Garcia, R., Vouimba, R. M., Baudry, M. & Thompson, R. F. The amygdala modulates prefrontal cortex activity relative to conditioned fear. Nature 402, 294–296, doi:10.1038/46286 (1999).

25 Gilmartin, M. R. & McEchron, M. D. Single neurons in the medial prefrontal cortex of the rat exhibit tonic and phasic coding during trace fear conditioning. Behav Neurosci 119, 1496–1510, doi:10.1037/0735-7044.119.6.1496 (2005).

26 Burgos-Robles, A., Vidal-Gonzalez, I., Santini, E. & Quirk, G. J. Consolidation of fear extinction requires NMDA receptor-dependent bursting in the ventromedial prefrontal cortex. Neuron 53, 871–880, doi:10.1016/j.neuron.2007.02.021 (2007).

27 Hoover, W. B. & Vertes, R. P. Anatomical analysis of afferent projections to the medial prefrontal cortex in the rat. Brain Struct Funct 212, 149–179, doi:10.1007/s00429-007-0150-4 (2007).

28 Vertes, R. P. Differential projections of the infralimbic and prelimbic cortex in the rat. Synapse 51, 32–58, doi:10.1002/syn.10279 (2004).

29 Heidbreder, C. A. & Groenewegen, H. J. The medial prefrontal cortex in the rat: evidence for a dorso-ventral distinction based upon functional and anatomical characteristics. Neurosci Biobehav Rev 27, 555–579, doi:10.1016/j.neubiorev.2003.09.003 (2003).

30 Franklin, K. B. J. & Paxinos, G. The mouse brain in stereotaxic coordinates. (Academic Press, 1997).

31 Sierra-Mercado, D., Padilla-Coreano, N. & Quirk, G. J. Dissociable roles of prelimbic and infralimbic cortices, ventral hippocampus, and basolateral amygdala in the expression and extinction of conditioned fear. Neuropsychopharmacology 36, 529–538, doi:10.1038/npp.2010.184 (2011).

32 Sørensen, A. T. et al. A robust activity marking system for exploring active neuronal ensembles. eLife 5, doi:10.7554/eLife.13918 (2016).

33 Stujenske, J. M. et al. Prelimbic cortex drives discrimination of non-aversion via amygdala somatostatin interneurons. Neuron 110, 2258–2267 e2211, doi:10.1016/j.neuron.2022.03.020 (2022).

34 Senn, V. et al. Long-range connectivity defines behavioral specificity of amygdala neurons. Neuron 81, 428–437, doi:10.1016/j.neuron.2013.11.006 (2014).

35 Klavir, O., Prigge, M., Sarel, A., Paz, R. & Yizhar, O. Manipulating fear associations via optogenetic modulation of amygdala inputs to prefrontal cortex. Nat Neurosci 20, 836–844, doi:10.1038/nn.4523 (2017).

36 Anastasiades, P. G. & Carter, A. G. Circuit organization of the rodent medial prefrontal cortex. Trends Neurosci 44, 550–563, doi:10.1016/j.tins.2021.03.006 (2021).

37 Little, J. P. & Carter, A. G. Synaptic mechanisms underlying strong reciprocal connectivity between the medial prefrontal cortex and basolateral amygdala. J Neurosci 33, 15333–15342, doi:10.1523/JNEUROSCI.2385-13.2013 (2013).

38 Little, J. P. & Carter, A. G. Subcellular synaptic connectivity of layer 2 pyramidal neurons in the medial prefrontal cortex. J Neurosci 32, 12808–12819, doi:10.1523/JNEUROSCI.1616-12.2012 (2012).

39 McGarry, L. M. & Carter, A. G. Prefrontal Cortex Drives Distinct Projection Neurons in the Basolateral Amygdala. Cell Rep 21, 1426–1433, doi:10.1016/j.celrep.2017.10.046 (2017).

40 McGarry, L. M. & Carter, A. G. Inhibitory Gating of Basolateral Amygdala Inputs to the Prefrontal Cortex. J Neurosci 36, 9391–9406, doi:10.1523/JNEUROSCI.0874-16.2016 (2016).

41 Manoocheri, K. & Carter, A. G. Rostral and caudal basolateral amygdala engage distinct circuits in the prelimbic and infralimbic prefrontal cortex. Elife 11, doi:10.7554/eLife.82688 (2022).

42 Klapoetke, N. C. et al. Independent optical excitation of distinct neural populations. Nature methods 11, 338–346 (2014).

43 Wietek, J. et al. A bistable inhibitory optoGPCR for multiplexed optogenetic control of neural circuits. Nat Methods 21, 1275–1287, doi:10.1038/s41592-024-02285-8 (2024).

44 Josselyn, S. A. & Tonegawa, S. Memory engrams: Recalling the past and imagining the future. Science 367, 39, doi:10.1126/science.aaw4325 (2020).

45 Burgos-Robles, A., Vidal-Gonzalez, I. & Quirk, G. J. Sustained conditioned responses in prelimbic prefrontal neurons are correlated with fear expression and extinction failure. J Neurosci 29, 8474–8482, doi:10.1523/JNEUROSCI.0378-09.2009 (2009).

46 Reppucci, C. J. & Petrovich, G. D. Organization of connections between the amygdala, medial prefrontal cortex, and lateral hypothalamus: a single and double retrograde tracing study in rats. Brain Struct Funct 221, 2937–2962, doi:10.1007/s00429-015-1081-0 (2016).

47 Kim, J., Pignatelli, M., Xu, S., Itohara, S. & Tonegawa, S. Antagonistic negative and positive neurons of the basolateral amygdala. Nat Neurosci 19, 1636–1646, doi:10.1038/nn.4414 (2016).

48 Zhang, X., Kim, J. & Tonegawa, S. Amygdala Reward Neurons Form and Store Fear Extinction Memory. Neuron 105, 1077–1093 e1077, doi:10.1016/j.neuron.2019.12.025 (2020).

49 Hagihara, K. M. et al. Intercalated amygdala clusters orchestrate a switch in fear state. Nature 594, 403–407, doi:10.1038/s41586-021-03593-1 (2021).

50 Printz, Y. et al. Determinants of functional synaptic connectivity among amygdala-projecting prefrontal cortical neurons in male mice. Nat Commun 14, 1667, doi:10.1038/s41467-023-37318-x (2023).

51 Kenna, M., Marek, R. & Sah, P. Insights into the encoding of memories through the circuitry of fear. Current Opinion in Neurobiology 80, 102712, doi:10.1016/j.conb.2023.102712 (2023).

52 Cummings, K. A. & Clem, R. L. Prefrontal somatostatin interneurons encode fear memory. Nat Neurosci 23, 61–74, doi:10.1038/s41593-019-0552-7 (2020).

53 Takehara-Nishiuchi, K. Neurobiology of systems memory consolidation. European Journal of Neuroscience 54, 6850–6863, doi:10.1111/ejn.14694 (2020).

54 Quirk, G. J., Repa, C. & LeDoux, J. E. Fear conditioning enhances short-latency auditory responses of lateral amygdala neurons: parallel recordings in the freely behaving rat. Neuron 15, 1029–1039, doi:10.1016/0896-6273(95)90092-6 (1995).

55 Repa, J. C. et al. Two different lateral amygdala cell populations contribute to the initiation and storage of memory. Nat Neurosci 4, 724–731, doi:10.1038/89512 (2001).

56 Collins, D. R. & Pare, D. Differential fear conditioning induces reciprocal changes in the sensory responses of lateral amygdala neurons to the CS(+) and CS(-). Learn Mem 7, 97–103, doi:10.1101/lm.7.2.97 (2000).

57 d’Aquin, S., et al. Compartmentalized dendritic plasticity during associative learning. Science 376, eabf7052, doi:10.1126/science.abf7052 (2022).

58 Sah, P., Faber, E. S., Lopez De Armentia, M. & Power, J. The amygdaloid complex: anatomy and physiology. Physiol Rev 83, 803–834, doi:10.1152/physrev.00002.2003 (2003).

59 Pitkanen, A., Savander, V. & LeDoux, J. E. Organization of intra-amygdaloid circuitries in the rat: an emerging framework for understanding functions of the amygdala. Trends Neurosci 20, 517–523, doi:10.1016/s0166-2236(97)01125-9 (1997).

60 Bolkan, S. S. et al. Thalamic projections sustain prefrontal activity during working memory maintenance. Nat Neurosci 20, 987–996, doi:10.1038/nn.4568 (2017).

61 Schmitt, L. I. et al. Thalamic amplification of cortical connectivity sustains attentional control. Nature 545, 219–223, doi:10.1038/nature22073 (2017).

62 Ghandour, K. et al. Orchestrated ensemble activities constitute a hippocampal memory engram. Nat Commun 10, 2637, doi:10.1038/s41467-019-10683-2 (2019).

63 Rothschild, G., Eban, E. & Frank, L. M. A cortical-hippocampal-cortical loop of information processing during memory consolidation. Nat Neurosci 20, 251–259, doi:10.1038/nn.4457 (2017).

64 Popa, D., Duvarci, S., Popescu, A. T., Lena, C. & Pare, D. Coherent amygdalocortical theta promotes fear memory consolidation during paradoxical sleep. Proc Natl Acad Sci U S A 107, 6516–6519, doi:10.1073/pnas.0913016107 (2010).

65 Clawson, B. C. et al. Causal role for sleep-dependent reactivation of learning-activated sensory ensembles for fear memory consolidation. Nat Commun 12, 1200, doi:10.1038/s41467-021-21471-2 (2021).

